# Dazl regulates germ cell survival through a network of polyA-proximal mRNA interactions

**DOI:** 10.1101/273292

**Authors:** Leah L. Zagore, Thomas J. Sweet, Molly M. Hannigan, Sebastien M. Weyn-Vanhentenryck, Raul Jobava, Maria Hatzoglou, Chaolin Zhang, Donny D. Licatalosi

## Abstract

The RNA binding protein Dazl is essential for gametogenesis, but its direct *in vivo* functions, RNA targets, and the molecular basis for germ cell loss in *DAZL* null mice are unknown. Here, we mapped transcriptome-wide Dazl-RNA interactions *in vivo*, revealing Dazl binding to thousands of mRNAs via polyA-proximal 3’UTR interactions. In parallel, fluorescence activated cell sorting and RNA-Seq identified mRNAs sensitive to Dazl deletion in male germ cells. Despite binding a broad set of mRNAs, integrative analyses indicate that Dazl post-transcriptionally controls only a subset of its mRNA targets, namely those corresponding to a network of genes critical for germ cell proliferation and survival. Additionally, we provide evidence that polyA sequences have key roles in specifying Dazl-RNA interactions across the transcriptome. Altogether, our results reveal a mechanism for Dazl-RNA binding, and illustrate that Dazl functions as a master regulator of a post-transcriptional mRNA program essential for germ cell survival.

## Introduction

RNA binding proteins (RBPs) are potent post-transcriptional regulators of gene expression. In the nucleus, RBPs can alter pre-mRNA processing to generate mRNAs with different coding and non-coding sequences. In the cytoplasm, RBPs influence mRNA localization, translation, and stability, typically through interactions with 3’ untranslated regions (3’UTRs). Importantly, the combined regulatory impacts of RBPs on RNA stability and translation can greatly influence protein expression. Despite widespread recognition of the importance of RBPs in regulating networks of mRNAs in development, immunity, and disease (Scotti and Swanson, 2016), our understanding of how specific RBPs modulate transcriptional programs to control cell fate is extremely limited.

Regulation of mRNA processing and translation is especially relevant during spermatogenesis, the highly ordered process of postnatal male germ cell development that yields haploid spermatozoa (Licatalosi, 2016). The first few postnatal days are critical for the establishment of spermatogonial stem cells which are required for continued sperm production throughout life (Manku and Culty, 2015). Many genes critical for spermatogonial proliferation and differentiation have been identified, including genes encoding cell cycle checkpoint factors, regulators of the DNA damage response, transcription factors, and RBPs that affect mRNA export and stability (Costoya et al., 2004; Raverot et al., 2005; Hao et al., 2008; Takubo et al., 2008; Pan et al., 2009; Saga, 2010; Song and Wilkinson, 2014). However, transcriptional and post-transcriptional mechanisms that coordinately control the expression of gene networks essential for germ cell maintenance are not well understood.

The importance of RBPs in germ cell development is well illustrated by the DAZ (<u>d</u>eleted in <u>az</u>oospermia) family of RBPs. These proteins comprise a family of germ cell restricted RBPs (Daz, Dazl, and Boule) necessary for gametogenesis in worms, flies, mice, and humans (VanGompel and Xu, 2011). Their significance was first demonstrated in the 1990s, when *DAZ* was discovered in a region of the Y chromosome deleted in 10-15% of men with azoospermia (no sperm) (Reijo et al., 1995; Reijo et al., 1996). Deletion of *DAZL* in mice leads to a dramatic decrease in the number of surviving germ cells (Ruggiu et al., 1997; Lin and Page, 2005). Remarkably, transgenic expression of human *DAZL* or *DAZ* partially rescues the extensive germ cell loss in *DAZL* knockout (KO) mice (Vogel et al., 2002), indicating functional conservation of DAZ RBPs across species. Despite the clear biological importance of DAZ proteins many critical questions remain, including the identities of their direct RNA targets, how these RNAs are regulated, and why loss of this regulation results in germ cell defects. In this study, we provide answers to these long-standing questions.

Dazl’s cytoplasmic localization, co-sedimentation with polyribosomes, and association with polyadenylated (polyA+) RNA (Ruggiu et al., 1997; Tsui et al., 2000) suggest potential roles in regulating germ cell mRNA stability or translation. In addition, yeast two hybrid analysis of Dazl-interactors identified RBPs with cytoplasmic roles in mRNA regulation including Pum2, QK3, and the polyA-binding protein Pabpc1 (Moore et al., 2003; Fox et al., 2005; Collier et al., 2005). However, the scarcity and variable number of germ cells present in *DAZL* KO mice (Ruggiu et al., 1997; Schrans-Stassen et al., 2001; Saunders et al., 2003) have presented major barriers to investigating Dazl’s direct *in vivo* function(s) in the male germline. Consequently, most previous Dazl studies have relied on reconstituted systems including transfection of somatic cell lines (Maegawa et al., 2002; Xu et al., 2013), artificial tethering to *in vitro* synthesized RNAs injected into oocytes or zebrafish (Collier et al., 2005; Takeda et al., 2009), and *in vitro*-derived germ cells (Haston et al., 2009; Chen et al., 2014) These efforts have suggested diverse functions for Dazl in different cell contexts, including roles in mRNA stabilization, stress granule assembly, mRNA localization, and translation (Fu et al., 2015). However, transfection assays have demonstrated that Dazl can have opposite effects on the same reporter RNA in different somatic cell lines (Xu et al., 2013). It is not clear whether these discrepancies are due to cell context and/or non-physiological levels or recruitment of Dazl to RNA targets.

While the direct *in vivo* functions are not defined, *in vitro* binding assays and X-ray crystallography of the Dazl RRM identified GUU as a high-affinity binding site (Jenkins et al., 2011). However, the frequency of GUU across the transcriptome hampers bioinformatic predictions of functional Dazl-binding sites and RNA targets *in vivo*. Microarray analyses of RNAs that co-immunoprecipitate (IP) with Dazl have suggested potential targets (Reynolds et al., 2005; Reynolds et al., 2007; Chen et al., 2014). Yet, few have been examined in *DAZL* KO mice, and these cannot account for the dramatic loss of germ cells observed. Furthermore, different investigators have arrived at alternate conclusions about Dazl’s role as a translation repressor or activator based on immunofluorescence (IF) assays of wild type (WT) and *DAZL* KO germ cells (Reynolds et al., 2005; Chen et al., 2014). Moreover, neither group explored whether the observed differences in protein abundance are associated with changes in mRNA levels.

Collectively, these observations underscore the need to identify Dazl’s direct *in vivo* RNA targets in an unbiased and transcriptome-wide manner, as well as new strategies to both isolate limiting *DAZL* KO germ cells for transcriptome-profiling and investigate how Dazl binds and regulates its RNA targets.

In this study, we used an integrative approach to elucidate the direct RNA targets and *in vivo* functions of Dazl in male germ cells. Multiple high-resolution, transcriptome-wide *in vivo* maps of Dazl-RNA interactions reveal Dazl binding to a vast set of mRNAs, predominantly through GUU sites in 3’UTRs. Using transgenic mice with fluorescently-labeled germ cells and FACS, we isolated germ cells from *DAZL* KO testes and wild type (WT) controls and used RNA-Seq to identify mRNAs that are sensitive to *DAZL*-deletion. Integrating the RNA-Seq and Dazl-RNA interaction datasets revealed that Dazl post-transcriptionally enhances expression of a network of genes with essential roles in spermatogenesis and cell cycle regulation. We also present multiple lines of evidence indicating that Dazl preferentially binds GUU sites in close proximity to polyA sequences, and demonstrate that the polyA tail at the 3’end of mRNAs has a critical role in Dazl-RNA binding. Collectively, these data provide important new insights into the mechanism of Dazl binding to its RNA targets, the molecular basis for postnatal germ cell loss caused by *DAZL* deletion, and reveal an mRNA regulatory program that is essential for postnatal germ cell survival.

## Results

### Dazl binds GUU-rich sequences across the testis transcriptome

To comprehensively map direct sites of Dazl-RNA interactions *in vivo*, HITS-CLIP libraries were generated from Dazl-RNA complexes purified from UV cross-linked adult mouse testes and sequenced using the Illumina platform (Figure 1A). The resulting CLIP reads from three biological replicates were filtered and mapped individually, and then intersected to reveal 11,297 genomic positions with overlapping CLIP reads in 3/3 libraries; hereafter designated as BR3 clusters (biologic reproducibility 3/3, Figure 1B). Remarkably, these interactions reveal Dazl directly binds over 3900 transcripts in adult testis (Supplemental File 1).

**Figure 1.**
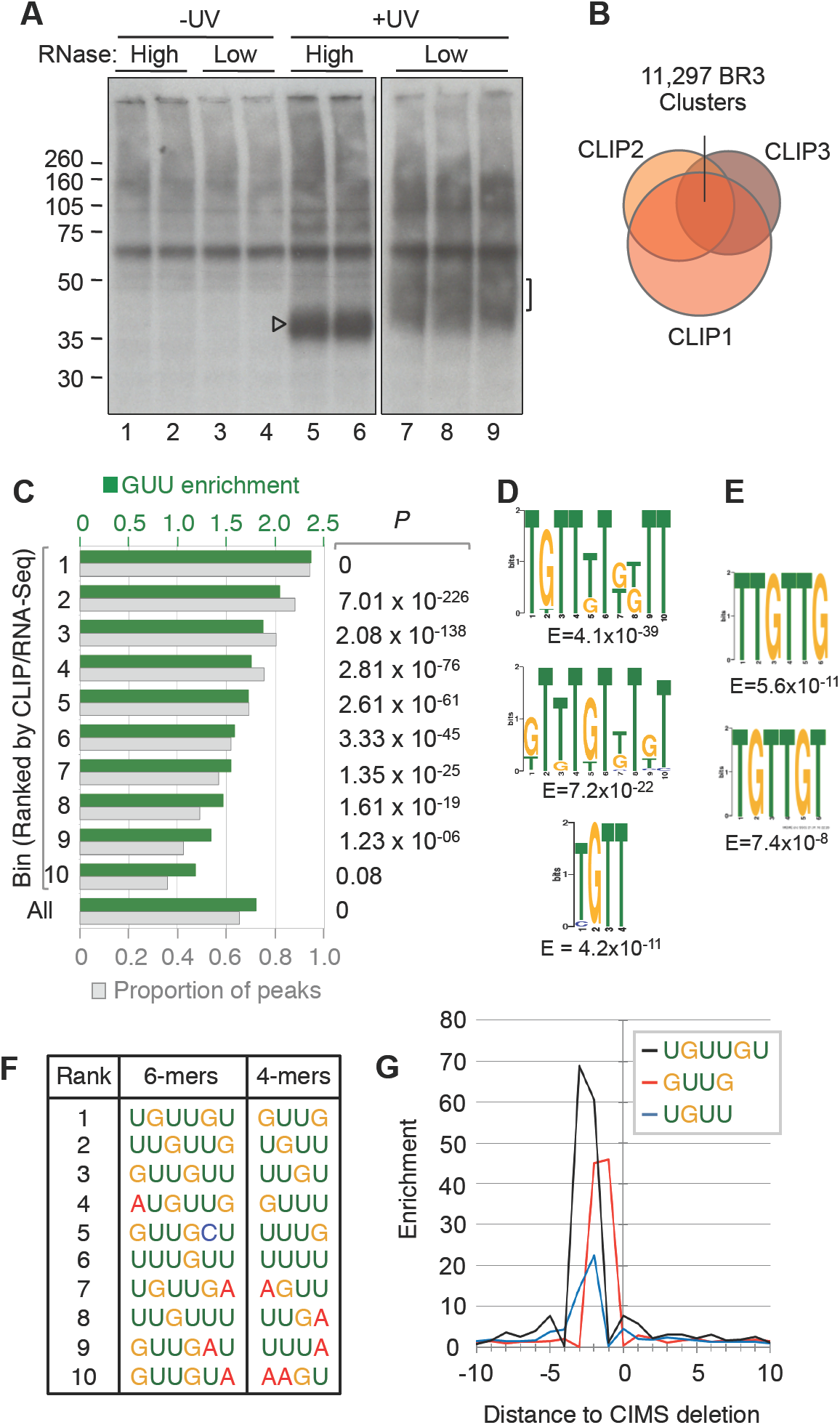
HITS-CLIP identification of Dazl-RNA contacts in the adult testis. **A.** Autoradiograph of nitrocellulose membrane containing radio-labeled cross-linked Dazl-RNA complexes purified from adult testes. Arrowhead indicates Dazl cross-linked to minimal RNA fragments. Bracket indicates region of nitrocellulose excised for cDNA preparation, which corresponds to Dazl cross-linked to RNA fragments of 35-50 nt **B.** Venn diagram showing the overlap between CLIP reads from each biologic replicate, and the identification of 11,297 genomic coordinates containing overlapping clusters of CLIP reads from 3/3 biologic replicates. **C.** Bins of CLIP peaks normalized to RNA-Seq, with GUU enrichment (green) and proportion of peaks in each bin that contain GUU (grey), with *P* values from χ^2^ distribution of GUU enrichment at right. **D.** Top motifs present in Bin 1, identified by MEME (top, middle) and DREME (bottom). **E.** Top motifs associated with CIMS sites identified by MEME. **F.** Top ten 6-mers and 4-mers associated with CIMS sites. **G.** Positioning of UGUUGU, GUUG, and UGUU motifs relative to CIMS deletion sites.

We next identified CLIP peaks within each cluster (Supplemental Figure 1). Consistent with X-ray crystallographic studies (Jenkins et al., 2011), these sites were enriched for GUU motifs compared to shuffled control sequences (1.8-fold, Figure 1C, All). Using RNA-Seq data from age-matched testes, CLIP peaks were normalized to RNA levels and parsed into 10 bins based on CLIP:RNA-Seq ratios. A clear correlation was evident between CLIP:RNA-Seq read ratio, GUU-enrichment, and the proportion of peaks in each bin that contained GUU (Figure 1C). Separately, *de novo* motif analysis using the MEME suite (Bailey et al., 2015) identified GTT-containing motifs as the most enriched sequence elements in genomic regions corresponding to peaks with the highest CLIP:RNA-Seq ratios (Figure 1D).

Dazl binding sites were also examined using an independent read mapping and analysis pipeline that takes advantage of cross-link induced mutation sites (CIMS) that reflect positions of protein-RNA cross-linking in HITS-CLIP data (Zhang and Darnell, 2011). *De novo* motif analysis of the top 1000 CIMS deletion sites (±10nt) identified GUU-containing motifs as the most enriched sequence elements (Figure 1E). Examination of 6mers and 4mers associated with the top 1000 CIMS sites ±20nt relative to background sequences identified enrichment of GUU-containing sequences around deletions, particularly the GUUG motif, which was enriched 40-60 fold around CIMS deletion sites, with cross-linking occurring at U residues within GUU triplets (Figure 1F,G). Thus, two independent read mapping and bioinformatic workflows confirm that Dazl predominantly binds GUU-rich sequences *in vivo*.

### Dazl predominantly binds GUU sites in close proximity to mRNA polyA tails

We next examined the distribution of Dazl-RNA contacts across the transcriptome and found that the majority of Dazl BR3 sites mapped to 3’UTRs of protein-coding genes (Figure. 2A; Supplemental Table 1). To improve the annotation of Dazl-3’UTR interactions we used PolyA-Seq to generate a quantitative global map of polyA site utilization in adult testes (Supplemental Figure 2A). Consistent with widespread transcription and alternative polyadenylation (APA) in the testis (Soumillon et al., 2013; Li et al., 2016), PolyA-Seq identified 28,032 polyA sites in RNAs from 16,431 genes (Figure 2B, Supplemental Figure 2B). Importantly, gene expression estimates from PolyA-Seq and RNA-Seq showed a strong correlation (R=0.83, Supplemental Figure 2C), as did estimates of polyA site usage from PolyA-Seq and qRT-PCR (R=0.72, 10 candidates examined; Supplemental Figure 2D).

**Figure 2.**
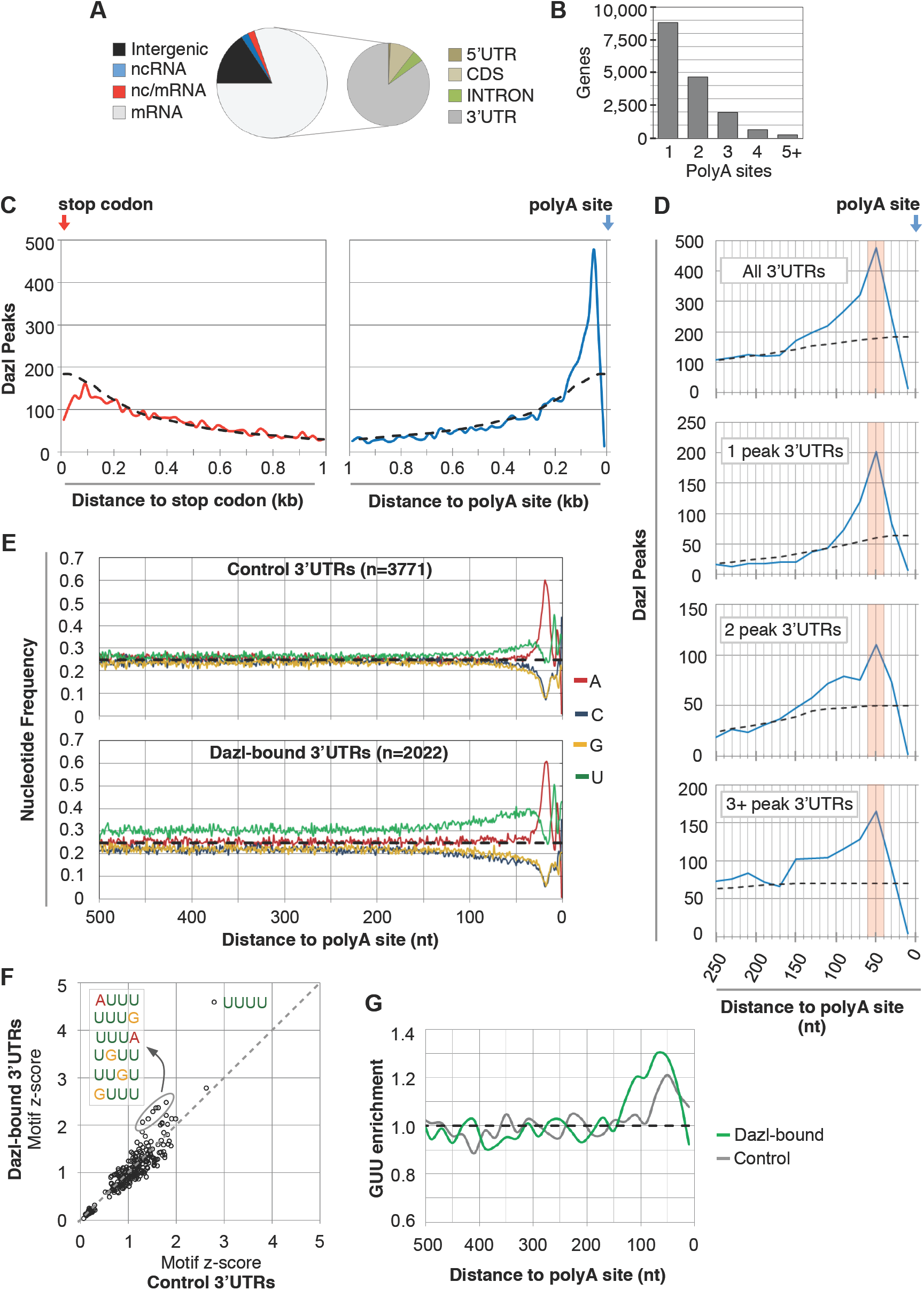
Dazl predominantly binds 3’UTRs at polyA-proximal sites. **A.** Distribution of BR3 CLIP sites in different genic regions, with the majority of interactions in mRNA 3’UTRs. BR3 clusters that mapped to regions with overlapping coding and non-coding RNAs were annotated as nc/mRNA. **B.** Number of genes with one or more polyA site, as determined by PolyA-seq analysis. **C.** Metagene analysis of the number of Dazl CLIP peaks in 20nt bins relative to the stop codon (left panel, red line) or the polyA site (right panel, blue line). Dashed line represents the expected random distribution. **D.** Top panel corresponds to a higher magnification view of the right panel in C. Bottom three panels show the distribution of CLIP peaks relative to the polyA site in 3’UTRs with one, two, and more than two peaks in the entire 3’UTR. **E.** Nucleotide frequency in the 500nt upstream of the polyA sites of control 3’UTRs (no Dazl CLIP reads, top panel), and 3’UTRs with Dazl-RNA interactions (bottom panel). **F.** Motif enrichment analysis of 3’UTRs with and without Dazl CLIP peaks (y and x axes, respectively), where values correspond to z-scores. **G.** GUU enrichment (observed/expected) in 3’UTRs of genes with and without Dazl CLIP peaks (green and grey lines, respectively).

Having defined 3’UTRs expressed in the testis, metagene analyses were performed to examine the distribution of 3’UTR Dazl-RNA interactions relative to the upstream stop codon and downstream polyA site. Compared to the expected distribution if peaks were randomly distributed (dashed line in Figure 2C), no positional preference was observed when Dazl-3’UTR interactions were measured relative to the stop codon (Figure 2C, compare red line to dashed line). In stark contrast, a prominent positional bias for Dazl-RNA contacts was observed relative to the polyA site, with strong enrichment within ∼150nt and the greatest number of interactions ∼50nt upstream of the polyA site (Figure 2C, compare blue line to dashed line). This positional preference is also evident on 3’UTRs with multiple Dazl CLIP peaks, with broadening of Dazl binding in a 3’-to-5’ direction as the number of peaks per 3’UTR increased (Figure 2D).

Further examination showed that 3’UTRs with Dazl BR3 CLIP sites had a higher proportion of uridine residues compared to a set of 3’UTRs lacking Dazl CLIP reads (Figure 2E, Supplemental Figure 3A). Moreover, the most enriched motifs in Dazl-bound versus control 3’UTRs were U-rich, with 2 of the top 7 most-enriched 4mers containing GUU (Figure 2F). Similarly, 3 of the top 4 6mers (and 7 of the top 20) contained GUU (Supplemental Figure 3B). Although both sets of 3’UTRs had enrichment of GUU upstream of polyA sites, GUU-enrichment was greater and extended over a broader region in Dazl-bound 3’UTRs (Figure 2G).

Collectively, transcriptome-wide mapping of direct, biologically-reproducible Dazl-RNA interactions in mouse testes indicates that Dazl binds U-rich 3’UTRs of thousands of genes expressed in the mouse testis, with preferential binding to GUU-containing sequences upstream of polyA sites.

### Dazl directly enhances steady state mRNA levels of a subset of in vivo targets

Significant barriers to understanding Dazl’s *in vivo* functions include the scarcity and variable number of germ cells in *DAZL* KO mice (Ruggiu et al., 1997; Lin and Page, 2005). To overcome these obstacles and collect *DAZL* KO germ cells for molecular analyses, we used Cre-lox to label germ cells with green fluorescent protein (GFP) followed by FACS. This approach (Zagore et al., 2015) utilizes the *Stra8-iCre* transgene that expresses Cre recombinase in postnatal germ cells (Sadate-Ngatchou et al., 2008), and the *IRG* transgene that expresses GFP following Cre-mediated recombination (De Gasperi et al., 2008). To generate animals with GFP+ Dazl-deficient postnatal germ cells (Figure 3A), we used mixed background (CD1xC57BL/6J) breeders heterozygous for the null *DAZL*^Tm1hgu^ allele (Ruggiu et al., 1997) (see Methods). Testes of *Stra8-iCre*^+^; *IRG*^+^; *DAZL*^*++*^ and *Stra8-iCre*^+^; *IRG*^+^; *DAZL*^Tm1hgu/Tm1hgu^ mice were indistinguishable at postnatal day 0 (P0), however germ cells declined steadily thereafter in the latter (data not shown). Cre expression from *Stra8-iCre* commences ∼P3, therefore we selected animals at P6 for tissue collection and FACS. At this age, spermatogonia are the only germ cells present. Importantly, germ cell-restricted GFP expression in *Stra8-iCre*^+^; *IRG*^+^; *DAZL*^*++*^ and *Stra8-iCre*^+^; *IRG*^+^; *DAZL*^Tm1hgu/Tm1hgu^ mice was confirmed by IF microscopy (Figure 3A), and germ cell markers were significantly enriched in GFP+ cells (data not shown). As expected, GFP+ spermatogonia were significantly reduced in *Stra8-iCre*^+^; *IRG*^+^; *DAZL*^Tm1hgu/Tm1hgu^ animals compared to littermate controls (Figure 3A-C).

**Figure 3.**
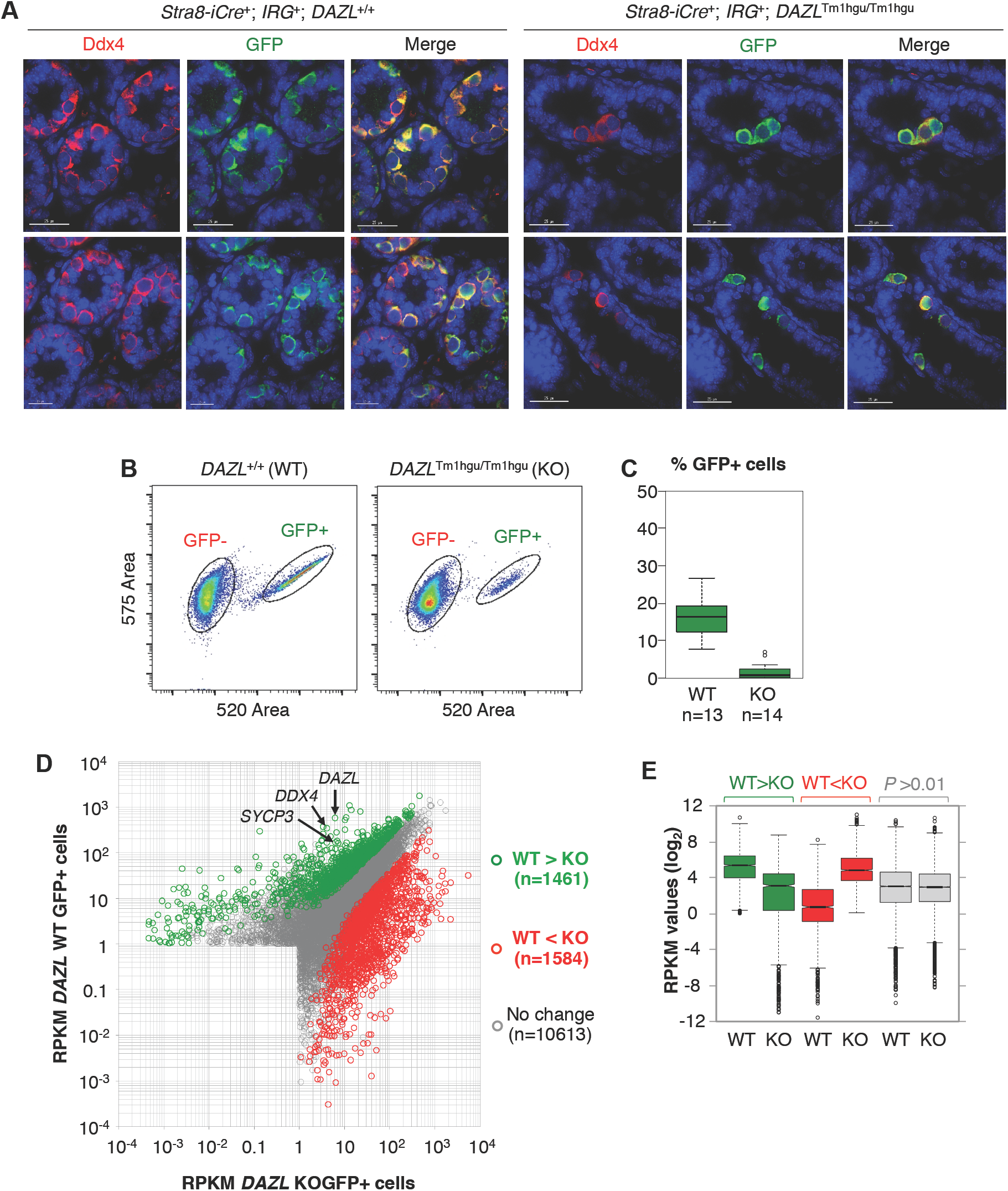
Genetic-labeling, isolation, and transcriptome-profiling of GFP+ WT and *DAZL* KO germ cells. **A.** Immunofluorescence analysis of Ddx4 and GFP expression in *Stra8-iCre*^+^; *IRG*^+^; *DAZL*^*++*^ (*DAZL* WT) and *Stra8-iCre*^+^; *IRG*^+^; *DAZL*^Tm1hgu/Tm1hgu^ (*DAZL* KO) testes at P6 (left and right panels, respectively). Sections from biological replicates are shown in the top and bottom rows. Scale bars represent 25μM. **B**. Flow cytometry plots of GFP- and GFP+ cells from WT and KO mice. **C**. Percentage of GFP+ cells isolated by FACS from P6 WT and *DAZL* KO mice. **D.** Scatter plot of RPKM values for genes in WT and KO cells. Green and red dots denote genes with decreased and increased RNA levels in Dazl KO cells compared to WT controls. **E.** Box plot showing the distribution of RPKM values for green, red, and grey genes indicated in panel D.

To determine how the absence of Dazl impacts global mRNA levels, RNA-Seq analysis was performed on GFP+ cells collected by FACS. Overall, the number of expressed genes and the distribution of their RPKM values were comparable between GFP+ WT and *DAZL* KO cells (11,739 and 12,269 expressed genes with RPKM>1, respectively). However, 1,462 transcripts had reduced RNA levels and 1,584 increased in *DAZL* KO cells (minimum 2-fold, adjusted *P*<0.01; Figure 3D, green and red dots, respectively; Figure 3E). Notably, *DDX4* and *SYCP3* were among the genes with reduced RNA in *DAZL* KO cells (86- and 26.5-fold, respectively; Figure 3D). Therefore, the reduced levels of *Ddx4* and *Sycp3* proteins previously observed in *DAZL* KO germ cells by IF and attributed to reduced translation (Reynolds et al., 2005; 2007) are associated with significant reductions in their corresponding mRNAs. Presently, we cannot exclude the possibility that differences in spermatogonial proliferation and differentiation contribute to RNA differences between P6 GFP+ WT and *DAZL* KO germ cells. However, markers of both undifferentiated (*CDH1*, *PLZF/ZBTB16*, and *GFRA1*) and differentiated spermatogonia (*KIT* and *STRA8*) had reduced RNA levels in *DAZL* KO cells.

To investigate which RNA changes are primary or secondary consequences of *DAZL* loss, we mapped Dazl-RNA interactions in P6 testes using iCLIP, which allows improved resolution of direct cross-link sites over HITS-CLIP. As with Dazl BR3 sites in adult testes, P6 BR3 regions were enriched for GUU sequences and the majority mapped to 3’UTRs (Figure 4A, B). To accurately map positions of Dazl-3’UTR contacts in spermatogonia, PolyA-Seq libraries were generated from GFP+ spermatogonia isolated by FACS from *Stra8-iCre*+;*IRG*+ testes. Of the 28,038 3’UTRs defined by PolyA-seq of whole adult testis (described above; Figure 2B), 16,502 were identified in the PolyA-seq data from GFP+ spermatogonia, corresponding to 10,370 genes. Intersecting the P6 iCLIP and spermatogonia PolyA-seq datasets showed that 84% of P6 Dazl BR3 sites (5,400/6,465) mapped to 3’UTRs of 2,290 genes, indicating Dazl directly binds to a vast network of RNAs in juvenile as well as adult germ cells via 3’UTR interactions. Furthermore, metagene analysis showed that Dazl-RNA interactions in spermatogonia-expressed 3’UTRs also had a polyA-proximal bias, with the greatest enrichment within 150nt of the polyA site (Supplemental Figure 4A). Thus, transcriptome-wide mapping of polyA sites and Dazl-RNA interactions confirm a polyA-proximal bias for Dazl-RNA binding to a broad set of mRNAs in both adult and juvenile testes.

**Figure 4.**
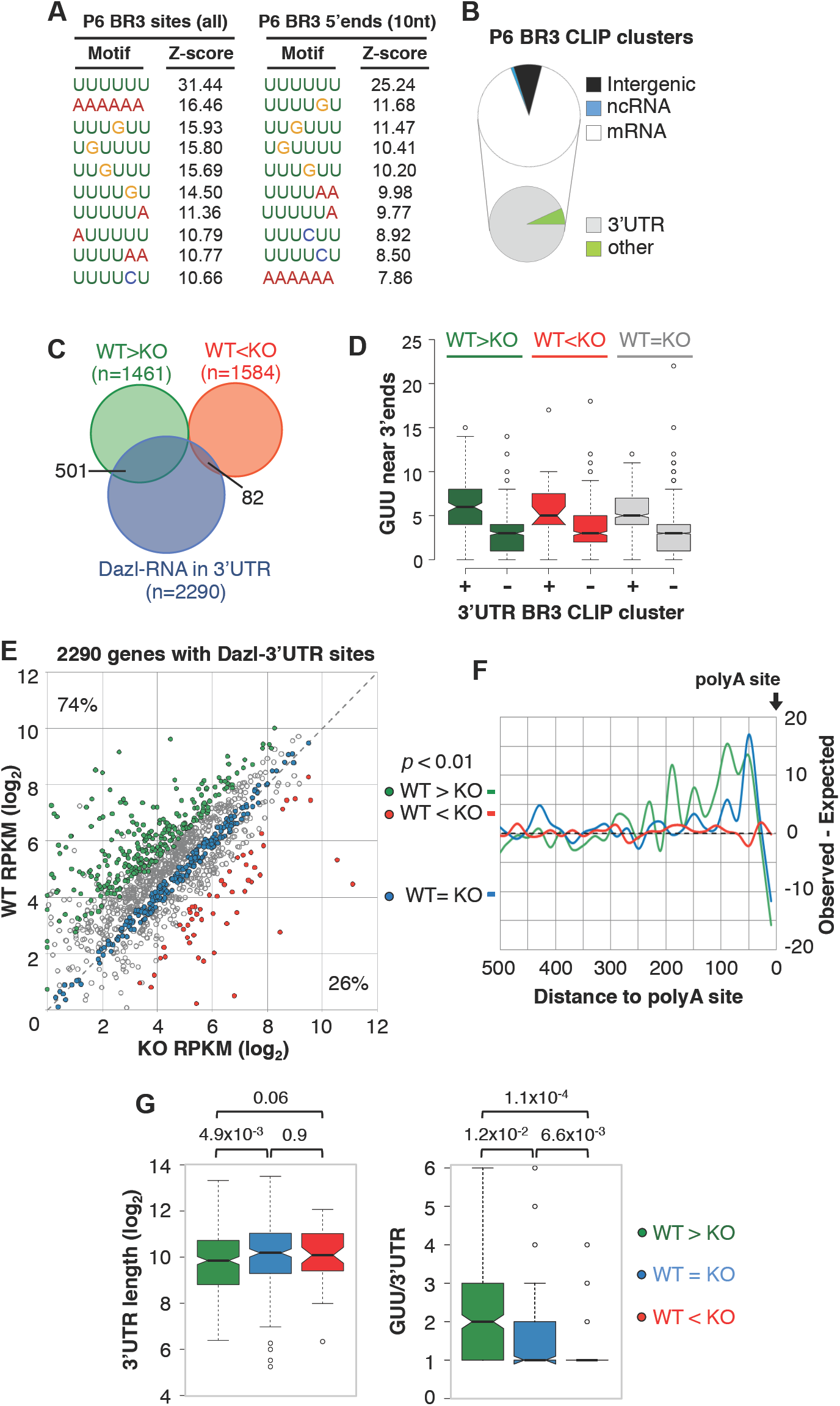
Dazl directly enhances mRNA levels through polyA-proximal binding. **A.** Top 10 motifs (based on z-score ranking) for sequences of BR3 regions (left) or 10 nucleotide windows centered at the 5’end of each BR3 CLIP region. **B.** Distribution of 6,465 BR3 Dazl-RNA sites from iCLIP analysis on P6 testes. **C.**. Venn diagram comparing genes with RNA level changes between WT and *DAZL* KO cells (2-fold or more, *P*<0.01) and the set of 2290 with Dazl-3’UTR interactions. **D.** Examination of GUUs in the last 250nt of 3’UTRs with and without BR3 sites identified by iCLIP analysis of P6 testes. **E.** RPKM levels of 2290 genes with Dazl-3’UTR interactions in GFP+ WT and GFP+ KO cells. Green, and red circles correspond to genes with RNA level differences between GFP+ WT and KO cells of 2-fold or more (adjusted *P*<0.01). Blue dots correspond to genes with RPKM values that do not differ more than 20% between WT and KO cells. **F**. Metagene analysis showing the distribution of BR3 Dazl-RNA sites in 3’UTRs of Dazl-enhanced, Dazl-repressed, and Dazl-insensitive genes (green, red, and blue, respectively). **G.** Examination of 3’UTR lengths (left) and frequency of GUU per 3’UTR (right) for genes with increased RNA, unchanged RNA, or decreased RNA leves (green, blue, and red, respectively) in GFP+ Dazl KO cells compared to WT controls. *P* values (Wilcoxon rank sum test) for pairwise comparisons indicated above.

Of the 2,290 genes with Dazl-3’UTR interactions in P6 testis, 583 also had differences in mRNA steady state levels in KO compared to WT germ cells. Strikingly, a disproportionate number of these overlapping genes (85%) had decreased RNA levels in KO cells (501 down-regulated and 82 up-regulated genes, respectively) (Figure 4C, E). To investigate why 3’UTR BR3 CLIP clusters are under-represented on transcripts that increase in *DAZL* KO cells, GUU frequencies were examined in 3’UTRs in different classes of genes with and without 3’UTR BR3 CLIP clusters. Independent of whether genes had increased, decreased, or unchanged RNA levels in KO compared to WT germ cells, 3’UTRs lacking BR3 sites had significantly fewer GUUs compared to 3’UTRs with BR3 sites (Figure 4D). The paucity of GUUs in 3’UTRs lacking BR3 CLIP clusters (in all gene categories) argues against ascertainment bias in the identification of RNAs directly bound by Dazl *in vivo*.

Metagene analyses revealed striking differences in the distribution of BR3 CLIP sites in the 3’UTRs of genes with increased or decreased RNA levels in KO versus WT cells. Whereas BR3 sites in 3’UTRs of *DAZL-*repressed genes showed no positional preference (Figure 4F, red line), Dazl binding sites in 3’UTRs of *DAZL*-enhanced genes were enriched near the polyA tail, with the greatest number of interactions within 100nt of the polyA site (Figure 4F, green line). Moreover, GUU motifs and Dazl BR3 interaction sites were significantly more abundant in *DAZL*-enhanced rather than *DAZL*-repressed genes, despite comparable 3’UTR lengths (Figure 4G). Taken together, these observations strongly suggest that Dazl’s directly regulated mRNA targets are those with reduced levels in KO germ cells, indicating that Dazl primarily functions as a positive post-transcriptional regulator of gene expression through polyA-proximal interactions.

We also examined 236 *DAZL*-insensitive genes with Dazl-3’UTR interactions but whose RNA levels did not differ by more than 20% between WT and *DAZL* KO cells (Figure 4E, blue dots). Remarkably, Dazl-RNA interactions in these 3’UTRs also displayed a strong polyA proximal bias (Figure 4F, blue line). Furthermore, we identified several examples where Dazl- 3’UTR binding patterns were indistinguishable between *DAZL*-insensitive and *DAZL*-enhanced 3’UTRs (Supplemental Figure 4B-D). These observations indicate that polyA-proximal binding alone is not sufficient for Dazl to enhance mRNA levels and suggest that Dazl acts in a 3’UTR specific manner. They also raise the possibility that a significant proportion of Dazl-3’UTR interactions may not have regulatory function and may represent opportunistic interactions reflecting the general mechanism of Dazl recruitment and/or stabilization on its RNA targets (see below).

### PolyA tracts are enriched near positions of Dazl-RNA interactions

Multiple lines of evidence suggest a potential role for polyA sequences in facilitating Dazl-GUU interactions. This includes the observed enrichment of Dazl-RNA contacts near polyA tails when metagene analyses were performed integrating CLIP and PolyA-Seq datasets from either adult or juvenile testes (Figure 2C, Supplemental Figure 4A). Additionally, AAAAAA was among the most enriched 6mers when regions of overlapping iCLIP reads were examined in BR3 clusters, but significantly less-enriched when motif analysis was performed on the 5’ends of iCLIP BR3 sites where cross-link sites are enriched (Konig et al., 2010) (Figures 4A, 6B). These observations indicate that genomic-encoded polyA sequences are enriched near, but not at, direct sites of Dazl-RNA cross-linking. Dazl was previously show to interact with Pabpc1 (Collier et al., 2005), which binds both polyA tails and genomic-encoded A-rich RNA sequences including auto-regulatory interactions in the 5’UTR of its own mRNA (Hornstein et al., 1999). Interestingly, *PABPC1* was among a small set of 47 genes with Dazl BR3 CLIP sites in the 5’UTR (56 sites in 5’UTRs of 47 genes, compared to 6,397 sites in 3’UTRs of 3,249 genes), prompting further examination of these outliers (Figure 5A; Supplemental File 1). For the majority (32/47 genes), CLIP read number was greatest in the 3’UTR and declined in a 3’-to-5’ manner (for example *ACYP1*, *CHIC2*, and *CCNI*, Figure 5B-D, 5’low genes). *PABPC1* was one of only 15 genes where more than 50% of the total CLIP reads were located in the 5’UTR (Figure 5E-F, 5’high genes). Marked differences were observed in the sequence features associated with these two sets of 5’UTRs. Consistent with low CLIP coverage, 5’UTRs of 5’low genes were not enriched for GUU-containing sequences. In contrast, 5’UTRs of 5’high genes resembled Dazl-bound 3’UTRs, whereby the most enriched motifs included polyU and polyU tracts interrupted by a single G (Figure 5G). Strikingly, AAAAA was among the most-enriched pentamers in 5’UTRs of 5’high genes, with a z-score equal to that of UGUUU (Figure 5G). Furthermore polyA tracts (5 A’s or more) were present in 73% in 5’high UTRs compared to 31% of 5’low UTRs (Supplemental Figure 5A). Thus, a common feature of biologically reproducible Dazl-RNA interactions across the transcriptome is the presence of local polyA sequence, either genomic-encoded or added to mRNA via 3’end processing.

**Figure 5.**
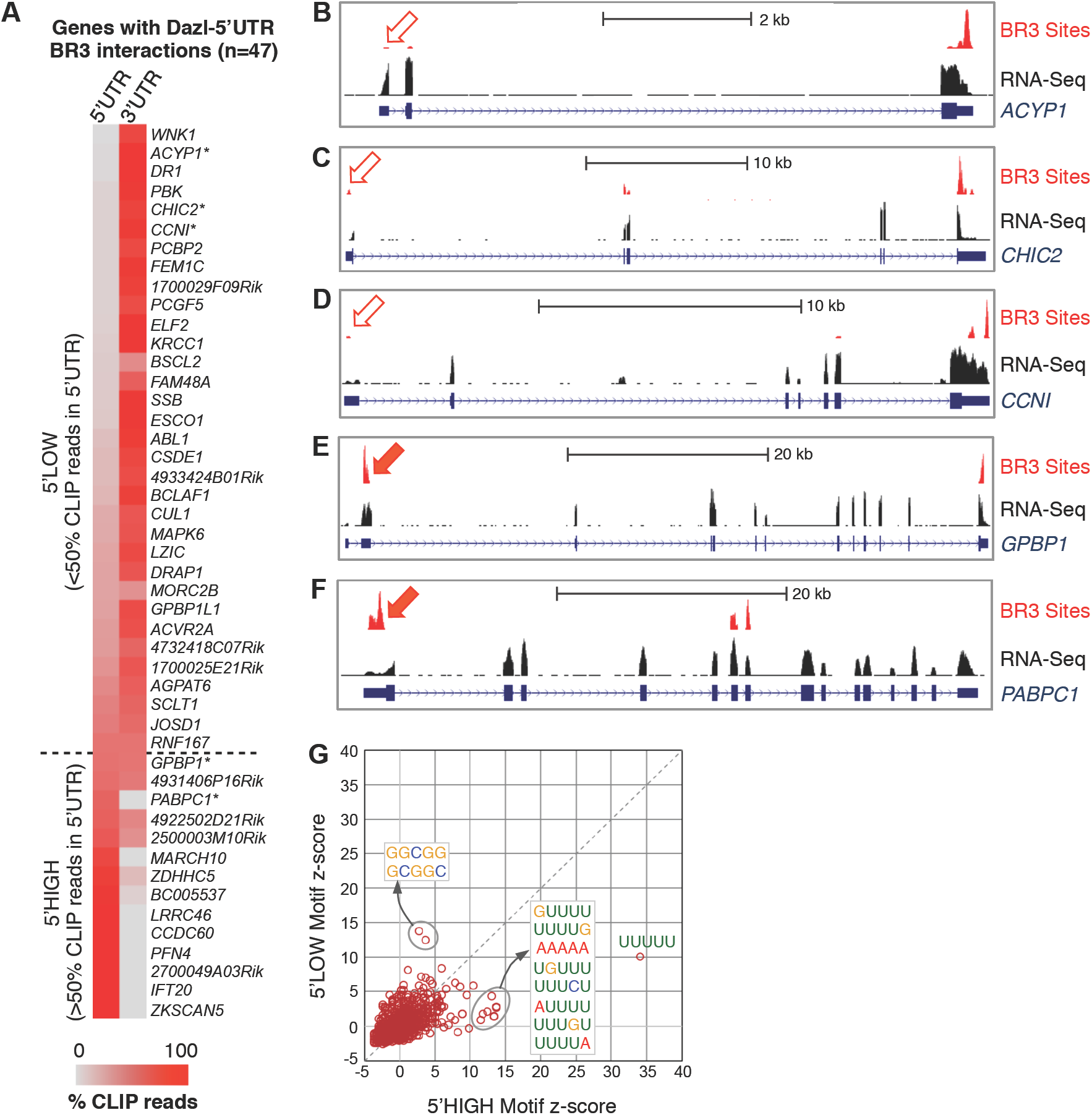
Rare Dazl-RNA interactions in 5’UTRs are associated with an enrichment of polyA tracts. **A.** Comparison of BR3 CLIP tag density in the 5’ and 3’UTRs of genes with 5’UTR BR3 CLIP sites. Color-coding represents the percentage of BR3 tags in the gene that map to either the 5’ or 3’UTR. Dashed line denotes cutoff between genes with less than (above) or greater than (below) 50% of the CLIP reads in the 5’UTR. **B-F.** Representative examples of genes listed in A, with RNA-Seq read density and CLIP read density in BR3 regions shown. **G.** Motif enrichment analysis of 5’UTRs with more or less than 50% of total BR3 CLIP reads in the 5’UTR.

### Dazl post-transcriptionally enhances gene expression in GC1-spg cells

Based on the data described above, we hypothesized that local polyA sequences facilitate Dazl-GUU interactions, thus providing a potential explanation for the preferential binding of Dazl to polyA-proximal GUU sites. To explore this possibility, we first established a stable monoclonal cell line that allows doxycycline (dox)-inducible expression of *DAZL*. GC-1sg cells were selected as the parental cell line, which are derived from spermatogonia (Hofmann et al., 1992) but express low levels of endogenous Dazl compared to mouse testis (Figure 6A). In these cells, dox-treatment results in predominantly diffuse cytoplasmic localization of Dazl, similar to the pattern observed in WT germ cells (data not shown). As in the testis, transcriptome-wide mapping in dox-treated GC1-sg cells showed that Dazl crosslink sites were significantly enriched for GUU-containing sequences (Figure 6B, Supplemental Figure 5B), with 80.7% of BR3 sites mapped to RefSeq mRNAs were in 3’UTRs (1226 genes, Figure 6C). Similar to P6 iCLIP data, AAAAAA was over-represented in entire BR3 regions with overlapping iCLIP reads (y-axis) but not at 5’ends (x-axis) where direct sites of Dazl-RNA cross-linking are enriched (Figure 6B).

**Figure 6.**
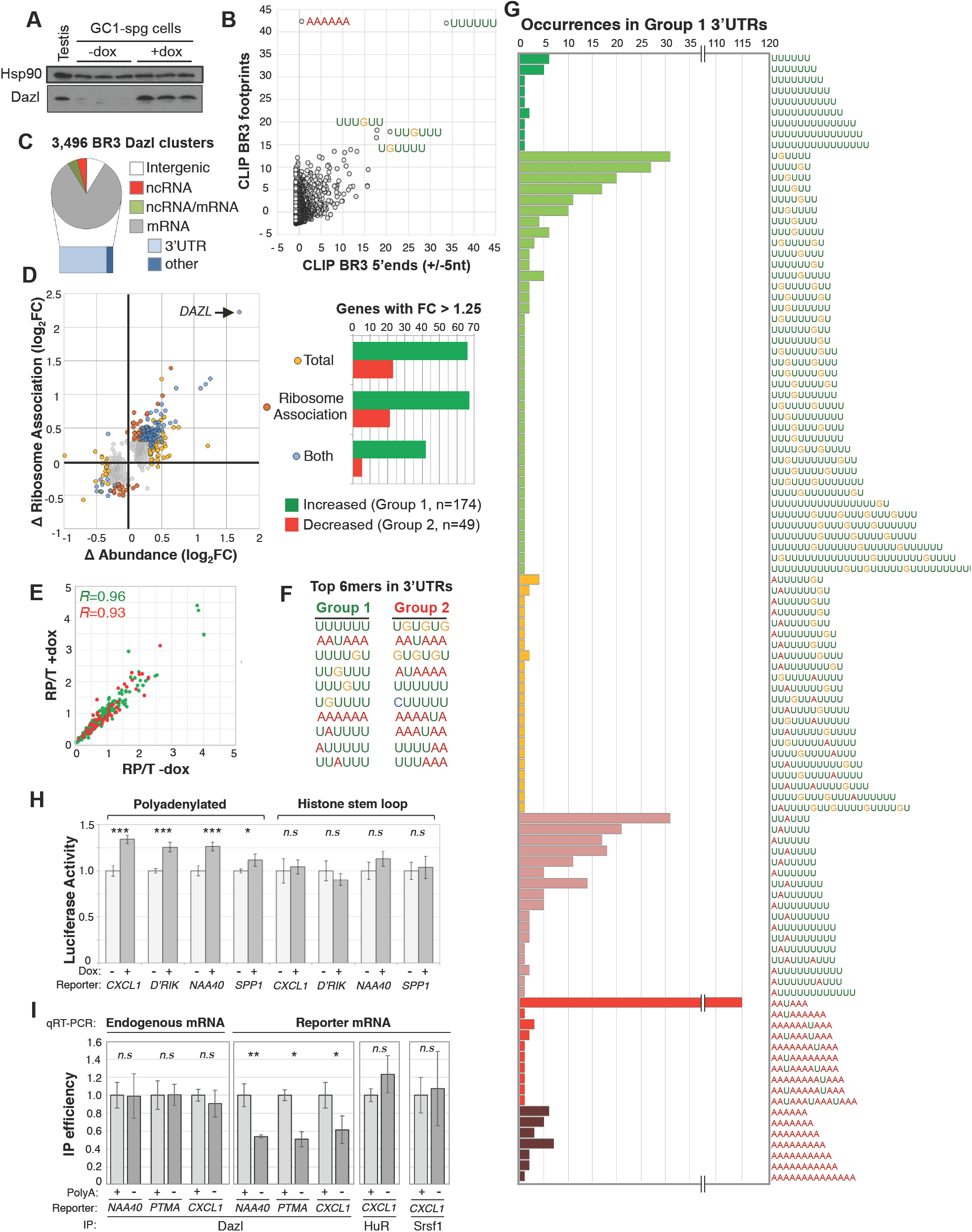
The polyA tail has an important role in Dazl-RNA binding and regulation. **A**. Western blot analysis showing dox-induced expression of *DAZL* in GC1-spg cells used for RNA-Seq analyses. **B**. Distribution of z-scores following motif-enrichment analysis of complete regions of BR3 positions bearing overlapping CLIP reads (y-axis) or 10 nt regions centered at the 5’end of BR3 sites (x-axis). **C**. Distribution of BR3 CLIP sites across the genome. **D**. Left: fold change in mRNA ribosome association (y-axis) and total mRNA level (x-axis) for 894 genes following dox-induction of *DAZL* expression. Right: Bar chart of 223 genes with 1.25-fold (or greater) change in mRNA association with ribosomes, change in total mRNA level, or both, following *DAZL* expression. **E**. Distribution of RP/T ratios for Group 1 and Group 2 genes (green and red, respectively) before and after dox-treatment. **F**. Top 10 most-enriched 6mers in last 250nt of Group 1 and 2 3’UTRs. **G**. Extended polyU tracts in Group 1 3’UTRs revealed by examining the distribution of the top 10 motifs (from F) in each Group 1 3’UTR and merging any that ovelap. **H**. Luciferase activity in untreated and dox-treated cells transfected with luciferase mRNA reporters bearing polyadenylated and non-polyadenylated 3’UTRs of the indicated genes. D’Rik corresponds to reporters bearing the 3’UTR of *D030056L22Rik*. **I**. Immunoprecipitation efficiency of endogenous *D030056L22Rik* mRNA and luciferase mRNA in dox-treated cells transfected with polyadenylated and non-polyadenylated versions of three different 3’UTRs. Values in H, and I are averages and standard deviations from 3 to 5 replicate experiments. *P*<0.0001***, *P*<0.01**, *P*<0.05*, *n.s* denotes no statistically significant difference.

To determine how *DAZL* induction affects mRNA abundance and translation in these cells, RNA-Seq datasets of total cytoplasmic RNA and ribosome-protected RNA fragments were generated from dox-treated and untreated cells. In total, 894 protein-coding genes had statistically significant differences in total cytoplasmic and/or ribosome-associated RNA levels between dox-treated and untreated cells (*P*<0.05, Figure 6D, left). Gene expression changes quantified by qRT-PCR and RNA-Seq had a strong positive correlation (R=0.9, 13 targets examined, Supplemental Figure 5C). The gene with the greatest mRNA differences was *DAZL*, corresponding to induction of the transgene, not the endogenous gene (evidenced by the absence of sequencing reads mapping to *DAZL*‘s 5’ and 3’UTRs, data not shown). Excluding *DAZL*, we focused on a set of 223 genes with mRNA level changes of 1.25-fold or greater for further analysis. The majority had increased cytoplasmic and/or ribosome-associated levels after dox treatment (174 dox-enhanced versus 49 dox-repressed genes) (Figure 6D, right). For both sets of genes, ribosome density was largely unchanged following dox-treatment, indicating that mRNA changes in ribosome-association following Dazl induction largely parallel changes in overall mRNA abundance (Figure 6E).

Comparing the 223 genes with dox-dependent RNA changes to those with Dazl-3’UTR interactions identified a common set of 44 genes, all of which belonged to the group of genes with dox-induced increased mRNA levels (total and/or ribosome associated). Similarly, when the 223 genes were compared to the 3907 with Dazl-3’UTR interactions in adult testes, the majority of the overlapping genes (91.7%) were enhanced by Dazl expression (67/174 enhanced versus 6/49 repressed). Consistent with these biased distributions, GUU was enriched in 3’UTRs of dox-enhanced genes, but underrepresented in 3’UTRs of dox-repressed genes (1.55-fold versus 0.85-fold, respectively). In addition, GUU-containing hexamers were among the most enriched motifs within 250nt of the polyA site of dox-enhanced genes (Figure 6F). Closer examination of the distribution of the top 10 most-enriched hexamers in dox-enhanced 3’UTRs showed that many overlap with one another to form extended polyU tracts interrupted by a single A, or more commonly, a single G (Figure 6G). Altogether, we conclude that genes with increased RNA levels following dox-treatment are Dazl’s direct targets in GC1-spg cells.

### Dazl binding and regulation of its RNA targets requires a polyA tail

Despite modest differences in RNA levels between untreated and dox-treated GC1-spg cells (compared to WT versus KO GFP+ spermatogonia), transcriptome profiling and CLIP analyses described above indicate that Dazl post-transcriptionally enhances a subset of its RNA substrates in GC1-spg cells via GUU-interactions in 3’UTRs, as in GFP+ spermatogonia. To test the hypothesis that the 3’ polyA tail is important for Dazl binding and RNA regulation, we generated a series of plasmids that express luciferase reporter mRNAs bearing the 3’UTR of Dazl target genes with either 1) the endogenous sequences necessary for 3’ end cleavage and polyadenylation, or 2) the stem loop structure from the 3’end of histone mRNAs to generate non-polyadenylated transcripts. Similar to effects on endogenous GC1-spg genes, Dazl-induction resulted in modest, yet highly reproducible increases in luciferase activity in cells transfected with polyadenylated reporters bearing the *CXCL1*, *D030056L22Rik NAA40*, and *SPP1* 3’UTRs (Figure 6H, left; Supplemental Figure 5D). In contrast, Dazl-induction did not result in increased luciferase levels in cells transfected with non-polyadenylated mRNAs (Figure 6H, right). We next performed co-immunoprecipitation (co-IP) assays to determine if these differences in luciferase levels were associated with differences in Dazl-3’UTR binding. While Dazl binding to endogenous mRNAs was unaltered in cells transfected with any of the reporter plasmids, Dazl binding was markedly reduced to reporter mRNAs bearing the histone stem loop compared to polyA+ versions (Figure 6I). This was true for reporters bearing the 3’UTRs from Dazl-enhanced genes (*CXCL1*, *NAA40*) as well as the *PTMA* 3’UTR which is bound by Dazl but insensitive to Dazl levels in testis and GC1-spg cells (Figure 6I, Supplemental Figures 4C,D, 5D). To determine if the histone stem loop causes broad remodeling of RBP-3’UTR interactions, we performed co-IP experiments to test binding of two RBPs (Srsf1 and HuR) previously shown to bind the *CXCL1* 3’UTR (Herjan et al., 2018). No difference in Srsf1 or HuR binding was observed to polyadenylated and non-polyadenylated luciferase reporters bearing the *CXCL1* 3’UTR (Figure 6I). Together, examination of multiple CLIP datasets and reporter mRNAs indicate that Dazl interaction with GUU-containing RNAs is facilitated by local polyA sequences. These observations provide a potential rationale for the widespread binding of Dazl to polyA-proximal GUU sites observed in thousands of RNAs in the germ cell transcriptome.

### Dazl post-transcriptionally controls a network of cell cycle regulatory genes

Collectively, the Dazl-RNA interaction maps, reporter mRNA assays, and RNA-Seq analyses of isolated germ cells and dox-treated cells in culture indicate that Dazl predominantly functions directly as a positive post-transcriptional regulator via 3’UTR interactions. Therefore, we more closely examined the 501 genes with Dazl-3’UTR interactions and reduced mRNA levels in GFP+ *DAZL* KO cells in order to identify potential primary causes of germ cell loss in *DAZL* KO mice. Hierarchical clustering revealed 17 groups of significantly enriched GO terms, with the majority (12/17) associated with different aspects of cell cycle regulation, including chromatin modification, condensation, or segregation; synaptonemal complex assembly; mitotic spindle checkpoint; DNA replication, repair, or packaging; and mitotic or meiotic cell cycle regulation (Figure 7A, groups with asterisks). The remaining groups contained genes and GO terms associated with RNA processing, ubiquitination, spermatogenesis, transcription by RNA pol II, and mRNA transport. Importantly, among the *DAZL*-enhanced genes are several that are necessary for spermatogenesis (Figure 7A,B, groups i, vi, and xiii), many with essential roles in spermatogonial maintenance or proliferation including *ATM* (Takubo et al., 2008), *DMRT1* (Zhang et al., 2016), *NXF2* (Pan et al., 2009), *SOHLH2* (Hao et al., 2008), *SOX3* (Raverot et al., 2005), *PLZF/ZBTB16* (Costoya et al., 2004), and *TAF4B* (Lovasco et al., 2015).

**Figure 7.**
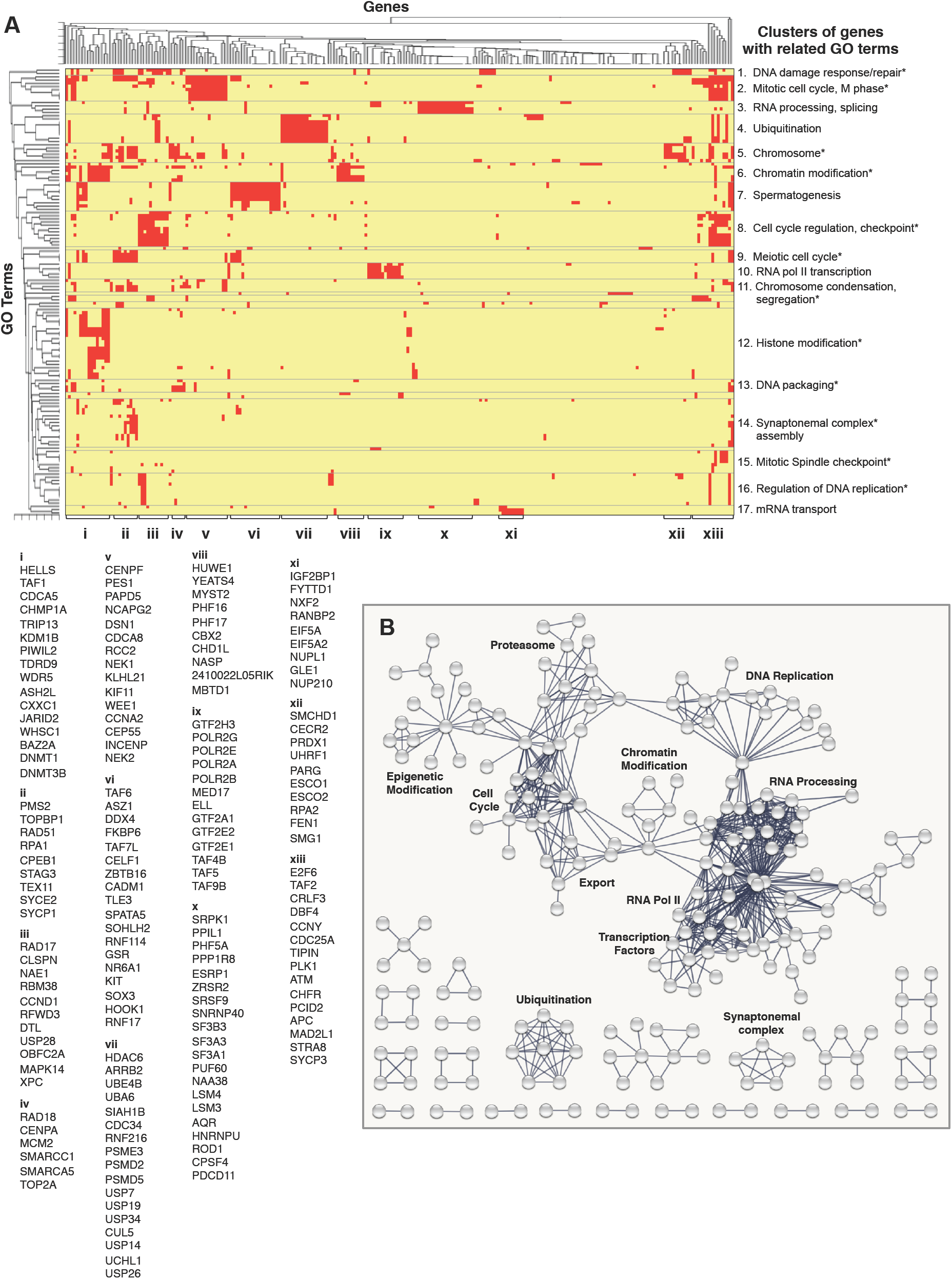
Dazl binds and regulates a broad network of essential genes encoding interacting proteins. **A.** Hierarchical clustering of enriched GO terms and genes associated with the set of 501 genes that are enhanced by Dazl and have Dazl-3’UTR interactions. Subgroups of overlapping GO categories are indicated at left, while subgroups of genes are indicated with letters at bottom. **B.** Lists of all genes in each letter group indicated at bottom of panel A. **C.** Protein-protein interaction network for genes with Dazl-3’UTR interactions and reduced RNA levels in Dazl KO cells compared to WT controls.

We also identified many examples in which genes encoding subunits of the same complex were among the 501 *DAZL*-enhanced genes (for example, gene cluster *ix* in Figure 7A,B). To further assess physical interactions between the protein products of the 501 genes, we used the STRING database of protein-protein interactions (PPIs) (Szklarczyk et al., 2015). Focusing on high confidence PPIs supported by experimental evidence or curated databases (minimum interaction score 0.9), we observed a higher than expected number of the 501 genes encode protein that interact with one another (PPI enrichment p-value = 0). This includes a broad network of interacting proteins including basal transcription factors, subunits of RNA pol II, RNA processing and export factors, chromatin binding proteins, epigenetic regulators, and several components of the proteasome and ubiquitination machineries (Figure 7B).

Together, GO and PPI enrichment analyses in combination with Dazl CLIP, PolyA-Seq, and RNA-Seq indicate that Dazl enhances postnatal germ cell survival via polyA-proximal interactions that promote high mRNA levels for a network of genes whose protein products physically interact, including several genes with essential roles in cell cycle regulation and germ cell survival.

## Discussion

The necessity of DAZ proteins for germ cell survival is well established in multiple species. However, the direct targets, regulatory roles, and biological functions of these RBPs remained unclear. Our integrative analyses combining transgenic mice, FACS, and a panel of unbiased, transcriptome-wide profiling tools provide important insights into the molecular and biological functions of this important family of RBPs.

### PolyA tracts facilitate Dazl-3’UTR interactions across the germ cell transcriptome

Transcriptome-wide mapping of direct Dazl-RNA interaction sites shows that Dazl binds thousands of mRNAs expressed in adult and juvenile testes, with most interactions in 3’UTRs. Given the widespread transcription and high RNA levels in male germ cells (Soumillon et al., 2013), it is not unexpected or uncommon for a germ cell-expressed RBP to reproducibly bind transcripts from thousands of genes (Hannigan et al., 2017). However, the Dazl CLIP maps are unusual in that the binding sites are highly enriched for GUU motifs near polyA tails at the 3’end of mRNAs, implying that additional sequence features specify which GUU sites are bound by Dazl. Our data suggest that widespread polyA-proximal Dazl-RNA interactions reflect a general mechanism of Dazl binding that depends on GUU and local polyA sequences *in vivo*. Supporting evidence includes the widespread preferential positioning of Dazl-RNA contacts upstream of polyA sites; over-representation of polyA tracts in the rare set of 5’UTRs with high CLIP read levels (including the *PABPC1* 5’UTR); enrichment of genomic-encoded polyA tracts downstream of direct sites of Dazl-RNA cross-linking in two independent iCLIP datasets; and failure of Dazl to efficiently bind and enhance reporter mRNAs lacking a polyA tail. Based on these observations and previous evidence of RNA-independent Dazl-Pabpc1 interactions (Collier et al., 2005), we propose that Dazl recruitment and/or stabilization on mRNA is mediated by local Pabpc1-polyA interactions. Considering the prevalence of GUU throughout the transcriptome, a requirement for local Pabpc1-polyA interactions in specifying sites of Dazl-RNA binding would increase the probability of Dazl loading on 3’UTRs, where the majority of cytoplasmic RBPs exert their regulatory functions (Sweet and Licatalosi, 2014).

### Dazl enhances mRNA levels of specific targets through polyA-proximal interactions

Regulation of mRNA translation and stability are intricately coupled processes (Bicknell and Ricci, 2017), therefore it is not clear whether Dazl functions primarily to prevent mRNA decay, or as a translational enhancer that indirectly impacts mRNA levels. It is also possible that Dazl’s primary function differs on distinct mRNAs. Although translation rates of specific mRNAs in WT and *DAZL* KO germ cells were not measured, we found that changes in mRNA steady state levels following *DAZL* expression in GC1-spg cells generally paralleled changes in mRNA association with ribosomes.

Additional studies are necessary to determine why some mRNAs with polyA-proximal interactions have reduced RNA levels in *DAZL* KO cells, while others are insensitive to *DAZL*-deletion. Notably, 3’UTRs of the former had the highest number of GUU sites per 3’UTR, despite modestly shorter lengths (Figure 4G). In addition, Dazl-RNA contacts on *DAZL*-insensitive genes were more concentrated near polyA sites, with reduced upstream 3’UTR binding compared to contacts in *DAZL*-enhanced genes (Figure 4F). Thus, multiple Dazl-RNA contacts may be necessary for Dazl to post-transcriptionally enhance expression of some genes. Intriguingly, examination of BR3 CLIP clusters with multiple distinct CLIP peaks showed that the peaks are often regularly spaced, with a median spacing of ∼60nt between adjacent peak regions (Supplemental Figure 1F), perhaps indicating Dazl homodimerization as seen by *in vitro* and yeast two-hybrid analyses (Ruggiu and Cooke, 2000). Binding to a broader segment of the 3’UTR in some mRNAs may displace or neutralize mRNA-specific *cis*- or *trans*-acting negative regulators that directly or indirectly promote mRNA decay or translational repression. However, we also identified several examples where the Dazl-RNA binding on 3’UTRs of *DAZL*-enhanced and *DAZL*-insensitive genes were indistinguishable suggesting that Dazl functions in a 3’UTR-specific manner (Supplemental Figure 4). These findings highlight the importance of combining RBP-RNA interaction maps with RNA-profiling to distinguish functional from potentially opportunistic RBP-RNA interactions.

### Dazl controls a network of genes essential for germ cell survival

Our RNA-Seq analysis of WT and *DAZL* KO germ cells shows a critical requirement for *DAZL* in maintaining the fidelity of the germ cell transcriptome through a combination of direct and indirect actions. While comparable numbers of genes had increased or decreased RNA levels in *DAZL* KO cells (1584 and 1461, respectively), they differed considerably with respect to enriched GO annotations and expression levels in WT cells. Genes with increased RNA in *DAZL* KO were significantly enriched for GO terms associated with the extracellular region and plasma membrane (Supplemental File 4). Strikingly, 40% of the genes in this set had little to no expression in WT cells (<1 RPKM). The presence of epigenetic regulators among the list of *DAZL*-enhanced genes with Dazl-3’UTR interactions (including *PIWIL2*, *TDRD9, DNMT1*, and *DNMT3B*, Figure 7A,B group i) may partially explain this ectopic gene expression, as well as the defects in erasure and re-establishment of DNA methylation observed in embryonic *DAZL* KO germ cells (Gill et al., 2011; Haston et al., 2009).

In stark contrast, genes with decreased RNA levels in *DAZL* KO cells were significantly enriched for GO terms associated with nuclear proteins involved in cell cycle regulation, DNA repair, DNA methylation, transcription, RNA processing, and RNA transport (Supplemental Figure 6, Supplemental File 4). Also enriched among *DAZL*-enhanced genes are GO terms associated with ubiquitin-protein ligase activity involved in cell cycle control, a broad set of genes with the annotation ‘spermatogenesis’, and numerous genes demonstrated to be essential for spermatogonial proliferation, differentiation, and/or survival (Supplemental Figure 6, group G). Representative examples of genes that are essential in spermatogonia and whose mRNA levels are significantly reduced in *DAZL* KO cells include *ATM* (Takubo et al., 2008), *CCNB1* (Tang et al., 2017), *DND1* (Yamaji et al., 2017), *HUWE1* (Bose et al., 2017), *KIT* (Blume-Jensen et al., 2000), *NXF2* (Pan et al., 2009), *PLZF/ZBTB16* (Costoya et al., 2004), *SOHLH2* (Hao et al., 2008) and *SOX3* (Raverot et al., 2005) (see Supplemental File 4 for a more extensive list). These observations suggest that Dazl functions as a regulator of regulators that directly sustains mRNA levels of essential genes through RNA binding, and indirectly by regulating mRNAs for transcription factors and epigenetic regulators that, in turn, define which genes are transcribed.

Global measurements of protein and mRNA stability have indicated that genes with short mRNA and protein half-lives are significantly enriched for the GO terms ‘transcription’, ‘cell cycle’, and ‘chromatin modification’ (Schwanhäusser et al., 2011), all of which are among the most-enriched GO terms associated with *DAZL*-enhanced genes with Dazl-3’UTR interactions. Strikingly, ∼50% of genes with reduced mRNA levels in *DAZL* KO cells are among the top 20% of genes with the highest mRNA levels in WT cells (Figure 3E). Conversely, genes with elevated mRNAs following *DAZL* expression in GC1-spg cells had very low levels of total and ribosome-associated mRNAs in the absence of dox, as well as significantly lower ribosome-density (Supplemental Figure 5E,F). Among these were *CXCL1* and *CXCL5*, two well-studied genes that are subject to extensive negative post-transcriptional regulation (Herjan et al., 2013). Furthermore, pre-mRNA levels of *DAZL*-enhanced genes were unaltered by Dazl expression (Supplemental Figure 5G), indicating that mRNA increases were due to regulation at the post-transcriptional level. Collectively, these observations suggest that Dazl functions in the germline to increase steady state levels of inherently unstable mRNAs to ensure high concentrations of regulatory factors important for spermatogonia proliferation.

In conclusion, our integrative analysis provides insights into the molecular basis of germ cell loss in *DAZL* KO mice and demonstrates that germ cell survival depends on a Dazl-dependent mRNA regulatory program. Given the functional conservation between mouse *DAZL*, human *DAZL*, and *DAZ* (Vogel et al., 2002), our study provides important insights into the molecular basis for azoospermia in 10-15% of infertile men with Y chromosome micro-deletions. Dazl’s RNA targets extend far beyond germ cell-specific genes and include many that encode core components of macromolecular complexes present in all proliferating cells. Therefore, our findings may also be relevant to other human diseases as *DAZL* is a susceptibility gene for human testicular cancer (Ruark et al., 2013), and is amplified or mutated in nearly 30% of breast cancer patient xenografts examined in a single study (Eirew et al., 2015). We propose a general model (Supplemental Figure 7) whereby Dazl binds a vast set of mRNAs via polyA-proximal interactions facilitated by Pabpc1-polyA binding and post-transcriptionally enhances expression of a subset of mRNAs, namely a network of genes essential for cell cycle regulation and mammalian germ cell maintenance. These observations provide new insights into molecular mechanisms by which a single RBP is recruited to its RNA targets and coordinately controls a network of mRNAs to ensure germ cell survival.

## Materials and Methods

### Animals and tissue collection

C57BL/6J animals bearing the *DAZL*^Tm1hgu^ allele were rederived at the Case Transgenic and Targeting Facility, and bred with animals bearing the *Stra8-iCre* or *IRG* transgenes. Mixed background (CD1xC57BL/6J) *Stra8-iCre*^++^; *DAZL*^Tm1hgu/*+*^ males and *IRG*^+^; *DAZL*^Tm1hgu/+^ females were crossed to generate *DAZL* WT and KO offspring with GFP+ male germ cells. Day of birth was considered P0. For all procedures, animals were anesthetized by isoflourane inhalation and death confirmed by decapitation or cervical dislocation. HITS-CLIP, iCLIP, adult testis RNA-Seq, and adult testis Poly-Seq were performed using CD-1 animals purchased from Charles River labs. Spermatogonia PolyA-Seq libraries were generated from FACS-isolated cells from 8 week *Stra8-iCre*^+^; *IRG*^+^ C57BL/6J males as previously described (Zagore et al., 2015). All animal procedures were approved by the Institutional Animal Care and Use Committee at CWRU.

### Cell isolation

Isolations of GFP+ cells was performed using dual fluorescence labeling as previously described (Zagore et al., 2015), except P6 cells were not stained with Hoechst and were collected using either BD Biosciences Aria or iCtye Reflection cytometry instrumentation.

### Microscopy

P6 testes were decapsulated in cold HBSS, fixed overnight in 4% paraformaldehyde, and paraffin-embedded by the Histology Core Facility at CWRU. Slides containing 5uM sections were de-paraffinized with 3 5-minute washes in xylene followed by a 5-minute rinse in 100% ethanol. Tissue was rehydrated with 5-minute incubations in 100%, 95%, 70% and 50% ethanol followed by tap water. Antigen retrieval was performed with citrate buffer, pH 6.0 (10mM Sodium Citrate, .05% Tween 20), at 95C for 20 minutes. Slides were cooled with tap water for 10 minutes followed by a 5-minute wash in 1X PBS. Tissue was permeabilized for 10 minutes in 0.25% TritonX-100 in 1X PBS and rinsed in PBST (0.1% Tween 20 in 1XPBS). Slides were blocked in 1% BSA in PBST for 1 hour. Sections were incubated with 1:500 anti-GFP (Abcam) and 1:500 anti-Ddx4 (Abcam) for 1 hour followed by 3 5-minute washes in PBST. An additional one hour incubation was performed using 1:100 anti-mouse Alexa-488 (ThermoFisher) and 1:100 anti-rabbit Cy3 (Jackson ImmunoResearch). Slides were washed 3 times in PBST before staining with 0.5ug/mL DAPI for 5 minutes and rinsing in 1X PBS. Tissue was mounted on coverslips with Fluoromount G (Southern Biotech) and images captured using a Deltavision Deconvolution Microscope.

### RNA-Seq

Adult testis RNA-Seq libraries (Tru-Seq, Illumina) were prepared from testes of two 8 week CD1 males. Illumina TruSeq/Clontech ultra low input library preparation with Nextera XT indexing was used to generate RNA-Seq libraries from GFP+ WT and *DAZL* KO cells purified by FACS. All RNA-Seq libraries were sequenced at CWRU Sequencing Core. Read mapping and gene expression quantification was performed using Olego and Quantas (Wu et al., 2013).

### PolyA-Seq

#### Sequencing library construction

Adult testis PolyA-Seq libraries were generated from 8 week CD1 mice. Spermatogonia PolyA-Seq libraries were generated from 400ng of RNA from spermatogonia isolated by FACS from 8 week old *Stra8-iCre*^+^; *IRG*^+^ males as previously described (Zagore et al., 2015). PolyA+ RNA was selected by oligodT-hybridization (Dynal) and fragmented by alkaline hydrolysis. RNA fragments 50-100 nt long were gel purified from 10% PAGE/Urea gels, and used as template for reverse transcriptions with SuperScript III (Invitrogen). First strand cDNA was gel purified, then circularized with CircLigase (EpiCentre). Circularized DNA was used as the template for PCR. After cycle number optimization to obtain the minimal amount of PCR product detectable by Sybr Gold (Invitrogen), the PCR product was gel purified and used as template for a second PCR with primers containing Illumina adaptor sequences. Adult and spermatogonia PolyA-Seq libraries were sequenced at CWRU and UC Riverside, respectively.

#### Read processing, filtering, and mapping

Reads consisting of two or more consecutive N’s or all poly(A) were filtered out, then remaining reads were processed to remove adapter and poly(A) sequences. Remaining reads were mapped to mouse genome mm9 with GSNAP allowing 2 mismatches. Reads with identical genomic footprint and 4N’s were then collapsed into single reads as likely PCR duplicates. Mapped reads that were likely internal priming events rather than true poly(A) tails were discarded if they had 6 or more consecutive A’s or 7 or more A’s in the 10 nt window downstream of the read 3’ end (cleavage and polyadenylation site). We accepted only 3’ ends with 1 of 14 hexamers within 50 nt upstream of the 3’ end (as described in (Martin et al., 2012).

#### Clustering of reads into poly(A) sites

To focus on high confidence poly(A) sites, we accepted read 3’ ends that were ≥ 1 tag per million reads (TPM). Due to heterogeneity in cleavage site selection, reads that fell within 10 nt of each other were clustered into poly(A) sites, and a TPM was calculated for the entire cluster (polyA site region).

#### Identification of 3’UTRs

PolyA sites regions from adult PolyA-Seq were intersected with RefSeq genes. Intergenic sites within 10kb of an upstream stop codon were assigned to the upstream gene. For each polyA site region mapped to a protein coding gene, the closest upstream stop codon was identified and used to define 28,032 3’UTRs in adult testes (only polyA sites with at least 5% of the total PolyA-Seq reads in a gene were considered). To identify 3’UTRs expressed in spermatogonia, mapped reads from spermatogonia PolyA-Seq libraries were intersected with 100nt regions corresponding to the 28,032 adult testis polyA sites, identifying 16,502 sites with read counts greater than zero in both spermatogonia PolyA-Seq replicate libraries.

#### qRT-PCR validation

qRT-PCR was performed as previously described (Zagore et al. 2015). Sequences of all primers used in this study are available in Supplemental File 6.

### Dazl HITS-CLIP from adult testes

#### Library construction

Dazl HITS-CLIP libraries were generated from testes from three biologic replicate 8-week old mice. Testes were detunicated in cold HBSS and seminiferous tubules UV-irradiated on ice. All steps were performed as previously described (Licatalosi et al., 2008; Licatalosi et al., 2012). Libraries were sequenced at the CWRU Sequencing Core and resulting reads mapped using Bowtie and Tophat (Trapnell et al., 2012) and filtered as previously described (Hannigan et al., 2017).

#### Identification of CLIP peaks

To identify peak regions in BR3 HITS-CLIP regions, the sum of all CLIP read lengths in a cluster was divided by the length of the cluster footprint to obtain an ‘expected’ density. Peaks are defined as regions of at least 10 nt long where the observed CLIP read density exceeded the expected density. 11,224 out of 11,297 had identifiable peaks (99.4% of BR3 clusters, 18364 peaks total).

#### Peak normalization and binning

For each BR3 peak, the sum of all observed-expected values per nt was calculated and divided by the average RNA-Seq read density per nt of the BR3 cluster. Only peaks within clusters with an average RNA-Seq read density per nt greater than 10 were considered for binning (13,106 peaks).

#### Genomic distribution of BR3 positions

To annotate BR3 sites, RefSeq coding regions, 5’UTRs, 3’UTRs, and introns were downloaded from the UCSC genome browser and intersected individually with BR3 coordinates.

#### Examination of genes with 5’UTR BR3 sites

For this analysis, only BR3 sites that mapped unambiguously to a single type of RefSeq gene fragment were examined (see Supplemental Figure 2).

### Dazl iCLIP from P6 testes and GC1-spg cells

#### Library construction and analysis

Individual Dazl iCLIP libraries were generated from three P6 testes. Three 10 cm plates of GC1-spg cells were induced with 10ng/mL doxycycline and harvested after 24 hours. The libraries were processed similar to HITS-CLIP libraries with the following changes. No 5’linker ligation step was performed, and the RT primer contained iSp18 spacers and phosphorylated 5’ end permitting circularization of first strand cDNA (after gel purification) to generate a PCR template without linearization (as described by (Ingolia, 2010). Libraries were sequenced at the CWRU Sequencing Core. Identical reads within each library were collapsed, and then mapped using Olego (Wu et al., 2013). Reads overlapping repetitive elements were discarded. After removing exon-spanning reads (0.06% of total), MultiIntersectBED (Quinlan and Hall, 2010) was used to identify genomic regions with overlapping CLIP reads from 3/3 libraries.

#### Genomic distribution of BR3 positions

BR3 regions were intersected with RefSeq genes and 3’UTRs of RefSeq protein coding genes.

### Examination of sequence features

#### Sequence analyses

Enriched motifs within CLIP regions and UTRs were performed using the EMBOSS tool Compseq. To generate z-scores, shuffled control sets were generated for each dataset analyzed using the EMBOSS tools Shuffleseq (10 shuffled versions of each sequence in each dataset). CIMS analysis was performed as previously described (Zhang and Darnell, 2011).

#### Distribution of Dazl-RNA contacts in 3’UTRs

Metagene analysis of Dazl-3’UTR interactions was performed on a subset of 3’UTRs defined by PolyA-Seq. 3’UTRs that overlapped with any intron sequence annotated in RefSeq were omitted. Only BR3 sites in genes with a single 3’UTR were analyzed. The resulting 5,284 BR3 HITS-CLIP peaks (adult testis) were examined in 2022 3’UTRs, and 3,321 BR3 iCLIP regions (P6 testis) were examined in 1,604 3’UTRs. To calculate distances, the midpoint of each BR3 site was measured relative to the upstream stop codon and the downstream polyA site. Positions of binding were then examined in 20nt windows relative to the stop codon or polyA site. For each 3’UTR, the expected distribution was determining by counting the number of BR3 sites in the 3’UTR and the number of 20nt bins in the 3’UTR, to determine the likelihood of any 20nt bin in a given 3’UTR containing a BR3 site. The control set of 3771 3’UTRs was identified by intersecting the 28032 3’UTRs from adult testis (identified by PolyA-Seq) with all adult testis CLIP reads, and selecting those with zero CLIP read coverage and no overlap with RefSeq introns.

### Generation of inducible stable cell lines

Stable monoclonal cell lines were generated using the T-Rex System (ThermoFisher) according to manufacturer’s instructions. The Dazl open reading frame was cloned into pcDNA 4/TO/myc-His C response element using the following primers: Dazl ORF F: (ACACTCGAGCCACCATGTCTGCCACAACTTCTGAGG), Dazl ORF R: ACAGGATCCTTAGCAGAGATGATCAGATTTAAGC)

### GC1-spg ribosome profiling and RNA-Seq

#### Library construction and analysis

Libraries were prepared using components of the following protocols with some modifications (Subtelny et al., 2014; Ingolia et al., 2012; Huppertz et al., 2014). Briefly, GC1-spg cells were induced with 10 ng/mL doxycycline and harvested after 24 hours. Cells were immediately scraped into lysis buffer containing the translation elongation inhibitor emetine. (10mM Tris-HCl(7.4), 5mM MgCl_2_, 100mM KCl, 1% TritonX-100, 0.1mg/mL emetine, 2mM DTT, cOmplete Mini EDTA-free protease inhibitor). 50% of the lysate was treated with 0.2U/uL RNaseI for 30 minutes at room temperature. The remaining lysate (gradient input) was reserved for RNA-Seq library preparation. Ribosome protected footprints were run on a 10-45% sucrose gradient and monosome associated RNA was purified and pooled. In parallel, RNA from the gradient inputs was fragmented by partial alkaline hydrolysis and size selected. Purified RNA fragments from monosome fractions and gradient inputs were DNase treated, rRNA depleted (RiboZero rRNA Removal kit – human/ mouse/ rat, Illumina) and processed concurrently. The libraries were prepared similar to iCLIP libraries except a preadenylated 3’ linker was used in place of the traditional 3’ RNA linker (Ingolia et al., 2012). Libraries were sequenced at the CWRU Sequencing Core. Identical reads within each library were collapsed and rRNA sequences removed. Read mapping, gene expression quantification, and differential gene expression analysis was performed using Olego and Quantas (Wu et al., 2013)

### Luciferase reporter generation

Dazl target 3’UTRs and at least 100nt of downstream sequence were cloned into the pRL-TK vector (Promega), replacing the SV40 late poly(A) region. To generate the non-polyadenylated reporter variants, the Hist2 histone stem loop (HSL) sequence was amplified from genomic CD1 mouse DNA using the following primers: Hist2 XbaI F (ACATCTAGAAAAAGGCTCTTTTCAGAGC), Hist2 BamHI R (TGTGGATCCTACCGTGACACAACTCTTTATC). After removing the SV40 late poly(A) region from pRL-TK, the HSL fragment was cloned downstream of the Renilla open reading frame. Dazl target 3’UTRs with disrupted poly A signals were inserted upstream of the HSL using blunt ended cloning.

### Dual Luciferase Assays

GC1-spg cells were induced with 1ng/mL doxycycline. After 24 hours, pRL-TK 3’UTR reporters and pGL4.54[luc2/TK] (Promega) firefly luciferase control plasmids were transfected into GC1-spg cells using Lipofectamine 2000 (Thermofisher). Media was replaced after 4-6 hours and cells were harvested after 24 hours. Dual luciferase assays were performed using the Dual-Luciferase Reporter Assay System (Promega) according to manufacturer’s instructions. Reporter renilla luciferase levels were normalized to firefly luciferase activity.

### Native RNA Immunoprecipitation

RIPs were preformed using conditions previously described (Gagliardi and Matarazzo, 2016) with the following antibodies: Dazl (Abcam ab34139), HuR LifeTechnologies (390600), CUG-BP1 Fisher (MA116675), SRSF1 Thermofisher (324500).

### GO analyses

GO term analyses were performed with the Cytoscape application BiNGO (Saito et al., 2012; Cline et al., 2007) using a hypergeometric statistical test and Benjamini & Hochberg FDR correction (significance level of 0.05) to identify enriched terms after multiple testing correction.. GO Slim settings were used to process the 656 set of genes (RPKM<1 in WT), while GO Full was used for the 1462 and 1584 sets of genes (Dazl-enhanced and repressed, respectively). A set of 13,659 genes with RPKM>1 in either WT or *DAZL* KO samples was used as the background gene set for enrichment. GO analysis of 501 Dazl-bound and enhanced genes was performed using GO Miner (Zeeberg et al., 2003).

### Protein-protein interactions

The STRING database (Szklarczyk et al., 2015) was used to identify PPIs between protein products of the 501 Dazl-enhanced genes with Dazl-3’UTR interactions. Selected parameters were experiments and databases (for interaction sources), and highest confidence (0.900) (for minimum required interaction score).

All deep sequencing datasets associated with this study are available at NCBI Gene Expression Omnibus accession number GSE108997.

## Author Contributions

L.L.Z. and D.D.L. conceived of and designed the study. L.L.Z. and T.J.S. performed the experiments. L.L.Z., T.J.S., M.M.H., S.M.W.V., C.Z. and D.D.L. analyzed the data. R.J. and M.H. provided reagents. L.L.Z. and D.D.L. wrote the manuscript.

## Acknowledgements

We are grateful to Marco Conti for providing *DAZL*^Tm1hgu/+^ mice, Jo Ann Wise for comments on the manuscript, and the following CWRU core facilities: Transgenic and Targeting Facility (mouse rederivation); Genomics (deep sequencing); Tissue Resources (embedding); Virology, Next Generation Sequencing, and Imaging Core (IF microscopy); Cytometry and Microscopy (FACS); Applied Functional Genomics (low input RNA-Seq libraries). This work was supported by funds from the National Institutes of Health to LLZ (T32 GM08056) and DDL (R01 GM107331).

## Supplemental Files

**Supplemental File 1. Adult testis HITS-CLIP BR3 clusters. Sheet 1** (‘11297 BR3 clusters’) contains genomic coordinates of 11,297 BR3 HITS-CLIP clusters (mm9) with CLIP reads per library, cluster length, and genomic annotation. **Sheet 2** (‘13106 binned BR3 peaks’) contains coordinates and values for normalized and binned peaks, as in Figure 1D, E. **Sheet 3** (‘47 genes 5’UTR BR3 clusters’) contains list of 47 genes with 5’UTR BR3 clusters, sorted by the percentage of total CLIP reads that map to the 5’UTR.

**Supplemental File 2. Adult testis PolyA-Seq and 3’UTR metagene data. Sheet 1** (‘28032 Adult Poly-Seq 3’UTRs’) contains genomic coordinates of 3’UTRs, corresponding gene, frequency of polyA site usage, and number of 3’UTRs per gene. **Sheet 2** (‘3907 genes Dazl-3’UTR’) contains list of genes with Dazl-RNA interactions in 3’UTRs defined by PolyA-Seq of adult testis. **Sheet 3** (‘Metagene’) contains 5284 CLIP peaks in 2022 3’UTRs used for metagene analysis, with coordinates of all features and distance of each peak midpoint to 3’UTR beginning and end.

**Supplemental File 3. Spermatogonia PolyA-Seq and 3’UTR metagene data. Sheet 1** (‘16502 SG Poly-Seq 3’UTRs’) contains genomic coordinates of 3’UTRs expressed in spermatogonia. **Sheet 2** (‘2290 genes Dazl-3’UTR’) contains list of genes with Dazl-RNA interactions in 3’UTRs defined by PolyA-Seq of spermatogonia. **Sheet 3** (‘Metagene’) contains 3321 P6 BR3 sites in 1604 3’UTRs used for metagene analysis, with coordinates of all features and distance of each BR3 midpoint to 3’UTR beginning and end. **Sheet 4** (P6 BR3 iCLIP sites) contains coordinates of P6 BR3 clusters, with number of CLIP reads per library and genomic annotation.

**Supplemental File 4. RNA-seq data for GFP+ cells. Sheet 1** (‘Gene expression’) contains RPKM values for all genes. **Sheet 2** (‘1462 genes decreased in KO’). **Sheet 3** (‘1584 genes increased in KO’). **Sheet 4** (‘656genes’) contains genes with average RPKM<1 in WT GFP+ cells. **Sheets 5** (‘307enhanced metagene’), **6** (‘236insensitive metagene’), and **7** (‘54repressed metagene’) contain BR3, 3’UTR, and distances for metagene analyses performed using P6 CLIP and spermatogonia PolyA-Seq data on *DAZL*-enhanced, insensitive, and repressed genes, respectively. **Sheet 8** (‘Essential Genes’) contains lists of *DAZL*-enhanced genes essential for mammalian germ cell development as determined by mutagenesis studies in mice. **Sheets 9, 10, and 11** contain GO results for 1462 *DAZL*-enhanced, 1584 *DAZL*-repressed, and 501 *DAZL*-enhanced genes with BR3-3’UTR interactions, respectively.

**Supplemental File 5. GC1-spg iCLIP data. Sheet 1**(GC1 BR3 iCLIP clusters) coordinates of BR3 clusters, with number of CLIP reads per library and genomic annotation. **Sheet 2** (‘RNA-Seq and ribosome profiling’) contains RPKM, fold change, and statistics for genes with changes in total and/or ribosome-associated RNA levels in dox-treated and untreated cells.

**Supplemental File 6.** Primers used in this study.

---

**Supplemental Figure 1.**
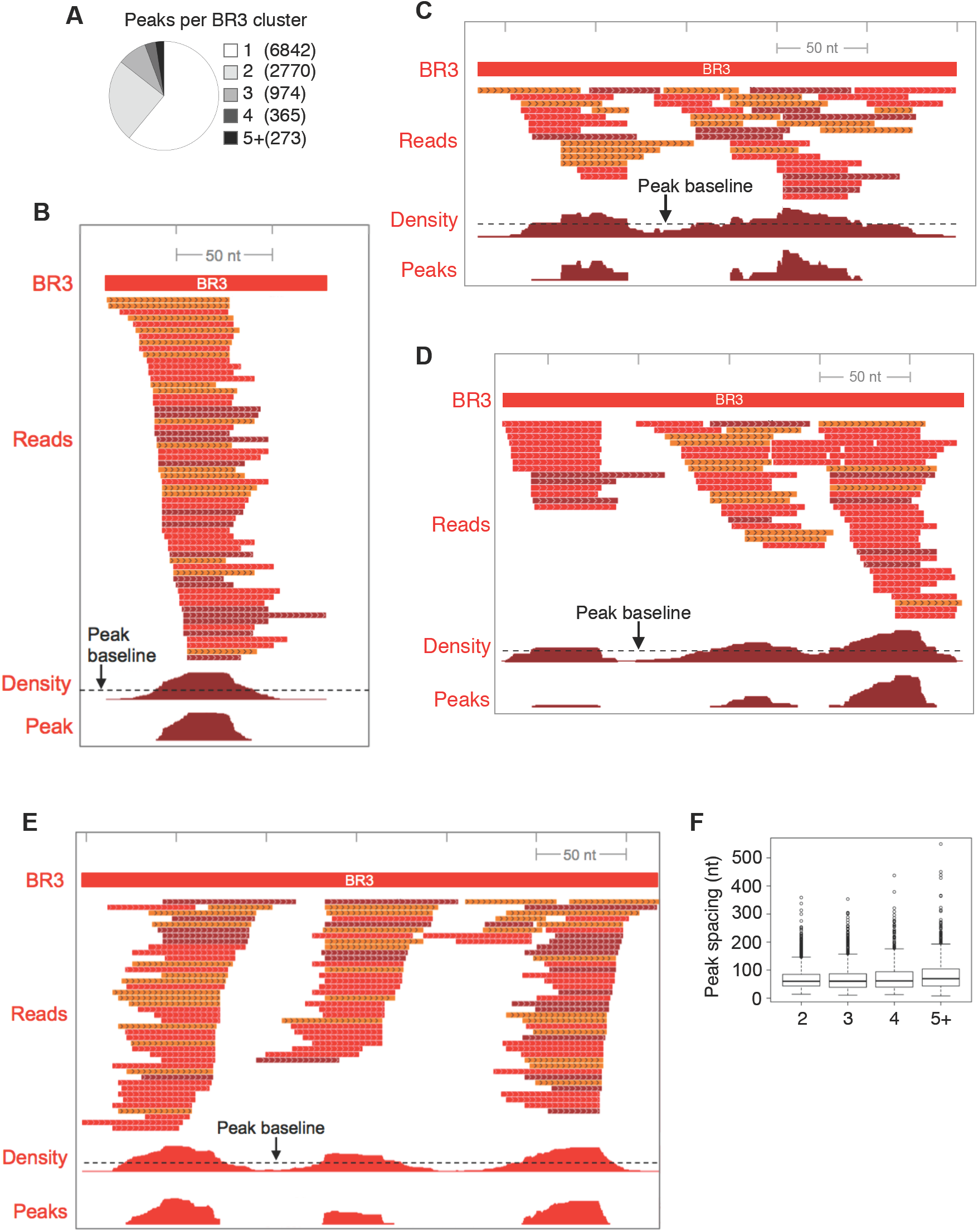
BR3 CLIP clusters with different numbers of peaks. **A**. Number of BR3 clusters with different numbers of peaks. BR3 clusters containing one (**B**), two (**C**), or three (**D**, **E**) peaks are shown. The footprint of the BR3 region is shown at the top. Reads are color coded to reflect each of the three CLIP libraries sequenced. Density indicates the number of overlapping reads per nucleotide within the BR3 region. Peak baseline corresponds to the total number of CLIP bases in the cluster divided by the length of the cluster. Peaks correspond to regions where the CLIP density exceeds the peak baseline. **F.** Distances between the midpoints of adjacent peaks in multi-peak clusters.

**Supplemental Figure 2.**
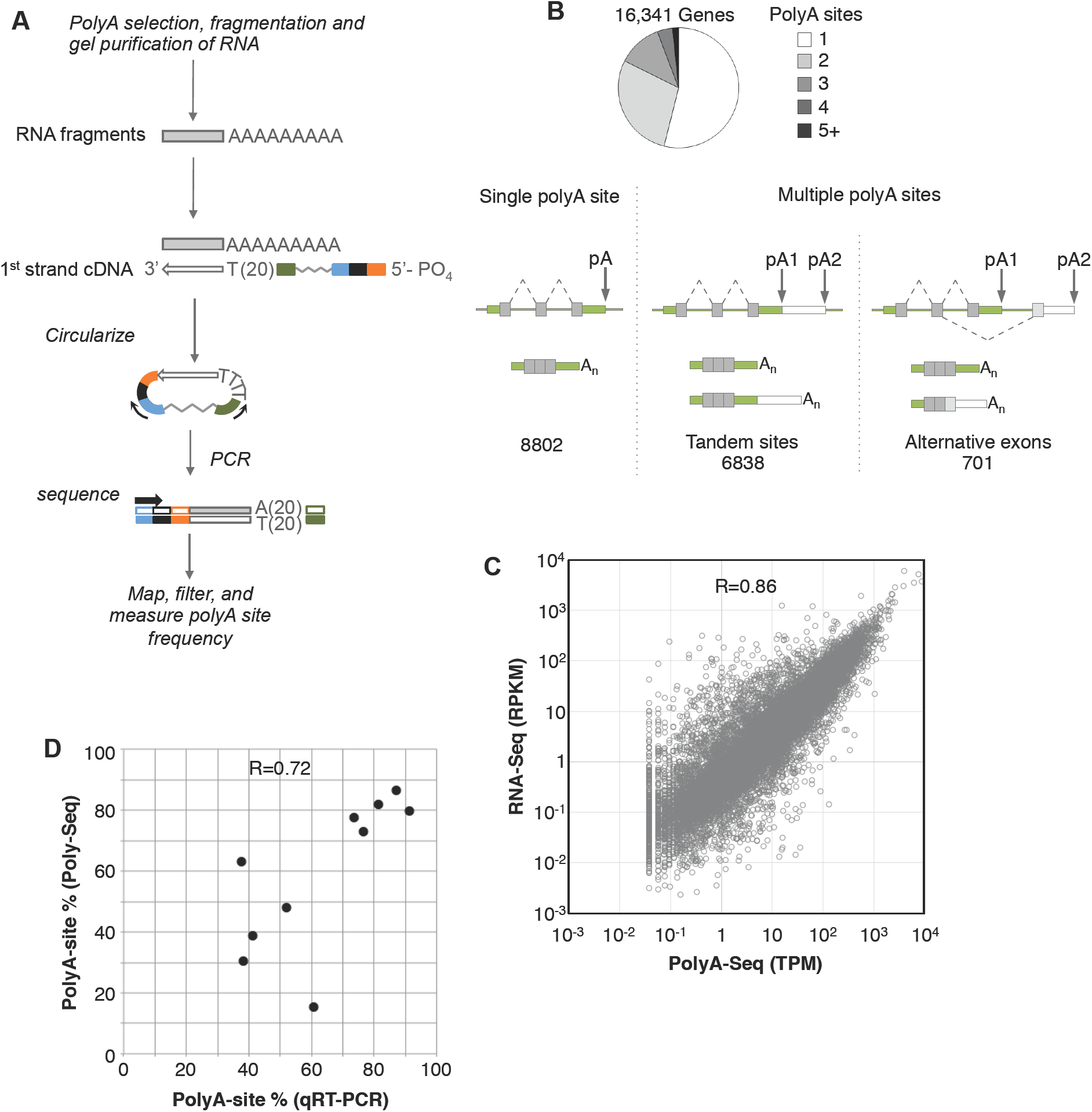
PolyA-seq identification of polyA sites of testis mRNAs. **A.** Overview of the PolyA-Seq library construction strategy. **B.** Characterization of polyA site usage in different genes. **C.** Comparison of RPKM and TPM values derived from RNA-Seq and PolyA-Seq, respectively, from adult testes. **D.** Comparison of proximal polyA site estimates from qRT-PCR and PolyA-Seq.

**Supplemental Figure 3.**
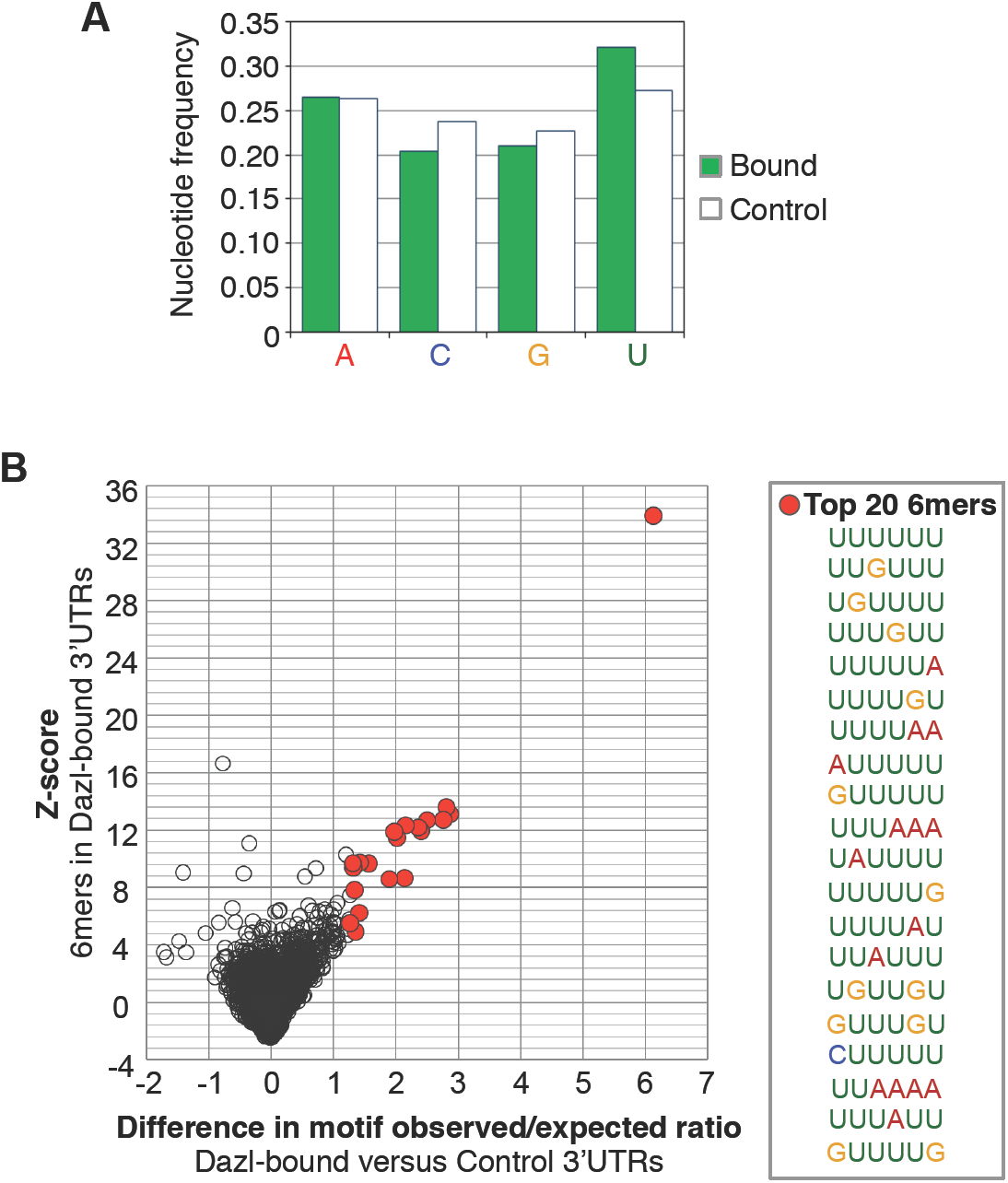
**A.** Nucleotide frequency in the last 500 nucleotides of 3’UTRs with Dazl BR3 sites (n=2022) and control 3’UTRs that have no CLIP tags (n=3771). B. Enrichment of 6-mers present in the 3’UTRs of genes with Dazl-3’UTRs compared to control genes. Each dot represents an individual 6 mer. X-axis corresponds to difference in observed-to-expected ratio for bound versus control 3’UTRs, while y-axis corresponds to the enrichment z-score for the 6-mer in the Dazl-bound 3’UTRs. Shown at right are the top 20 6-mers ranked by difference in the observed-to-expected ratios between bound and control 3’UTRs.

**Supplemental Figure 4.**
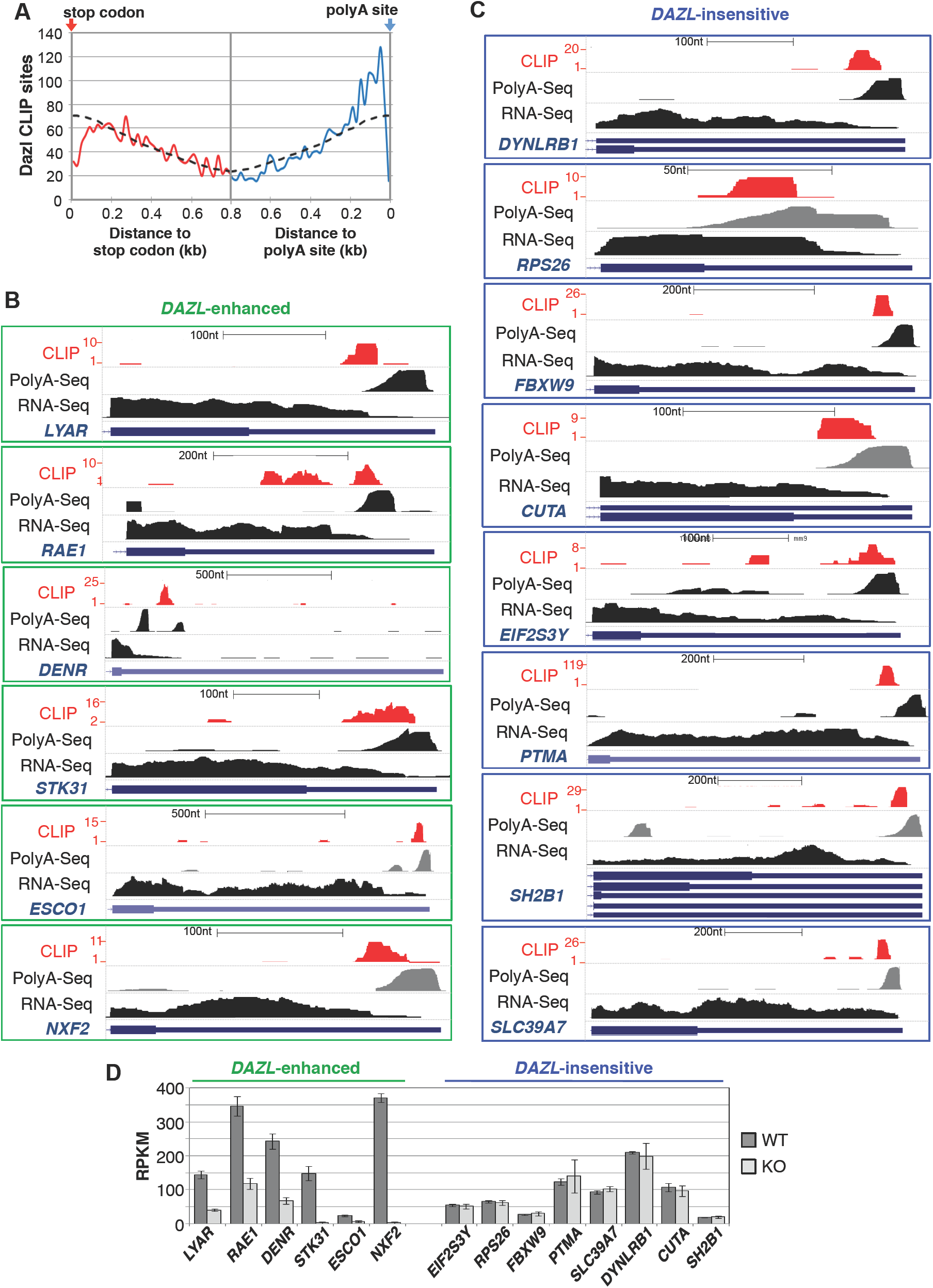
**A.** Metagene analysis of the distribution of BR3 Dazl-RNA sites (from P6 iCLIP) in 3’UTRs of spermatogonial-expressed 3’UTRs (as in Figure 2C). Genome browser images showing P6 BR3 CLIP sites in 3’UTRs of Dazl-enhanced and Dazl-insensitive genes (**B** and **C**, respectively). Adult PolyA-seq and RNA-Seq tracks are also shown. CLIP scale denotes the maximum number of overlapping CLIP tags in the region shown. **D**. Average RPKM values for the genes shown in panels A and B.

**Supplemental Figure 5.**
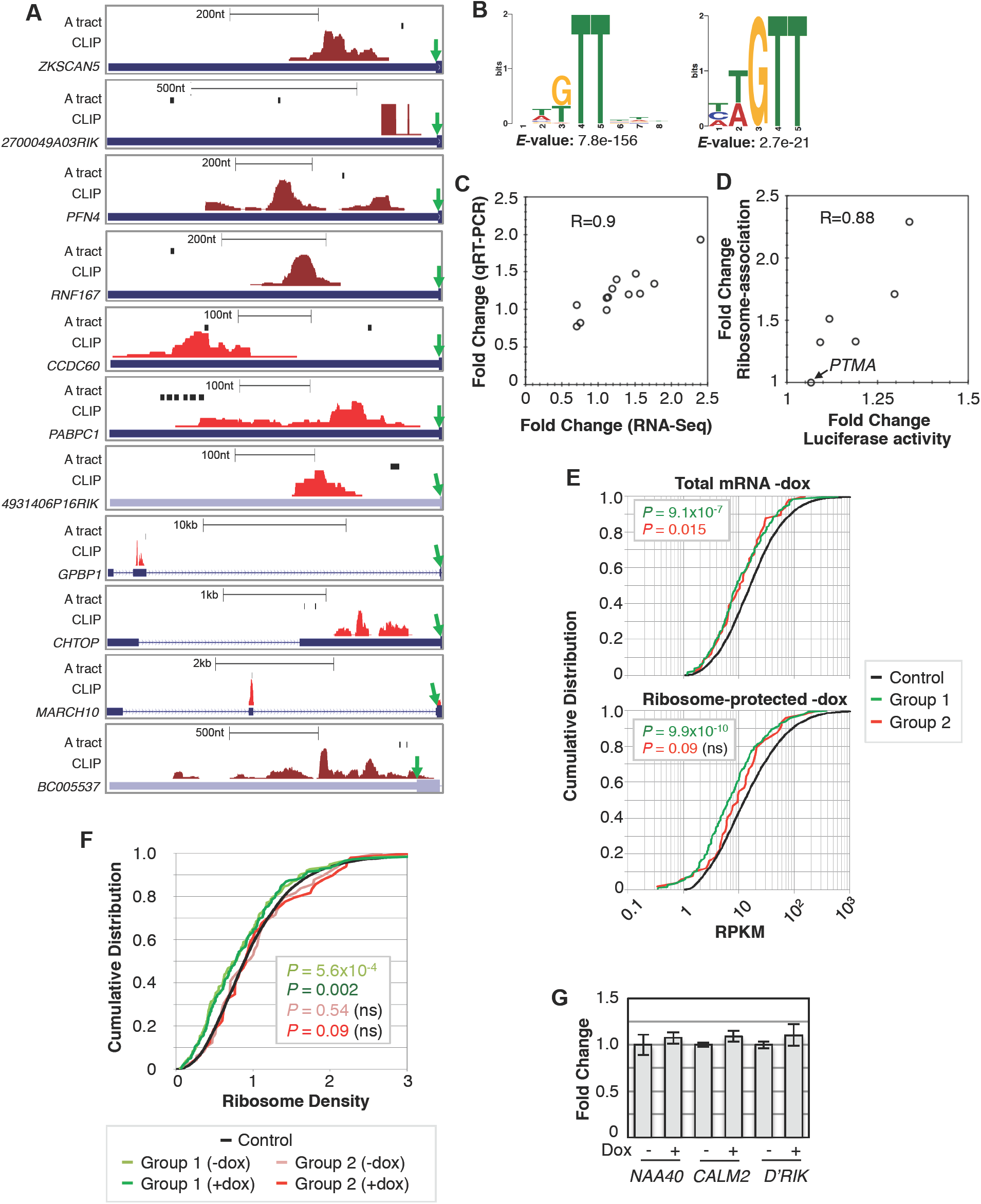
**A.** Genome browser images of 11 5’UTRs with Dazl BR3 CLIP clusters (adult testis) and polyA tracts of 5 A’s or more. PolyA tracts indicated by black boxes. Green arrows denote the start codon. **B.** The most significant motifs (identified by the programs MEME and DREME, left and right respectively) in 10nt regions centered at the 5’end of Dazl BR3 CLIP sites from GC1-spg cells. **C.** Comparison of dox-induced RNA level fold changes determined by qRT-PCR and RNA-Seq (13 genes represented). **D.** Comparison of dox-induced fold changes in luciferase activity for mRNA reporters bearing different 3’UTRs (x-axis) versus fold changes in ribosome-association for the corresponding endogenous transcripts determined by deep sequencing (y-axis). **E.** Cumulative distribution of RPKM values for control, Group 1 (enhanced), and Group 2 (repressed) genes (black, green, red lines, respectively) in untreated GC1-spg cells. *P* values (Wilcoxon rank sum test) correspond to pairwise comparisons of Group 1 (green) or Group 2 (red) RPKM values to control. **F.** Cumulative distribution of ribosome density values for Group 1 and 2 genes before and after dox-treatment. Ribosome density is the ratio of RPKM values from ribosome-protected fragments to total RNA for each gene. **G**. qRT-PCR analysis of pre-mRNA levels for three genes in dox treated and untreated cells.

**Supplemental Figure 6.**
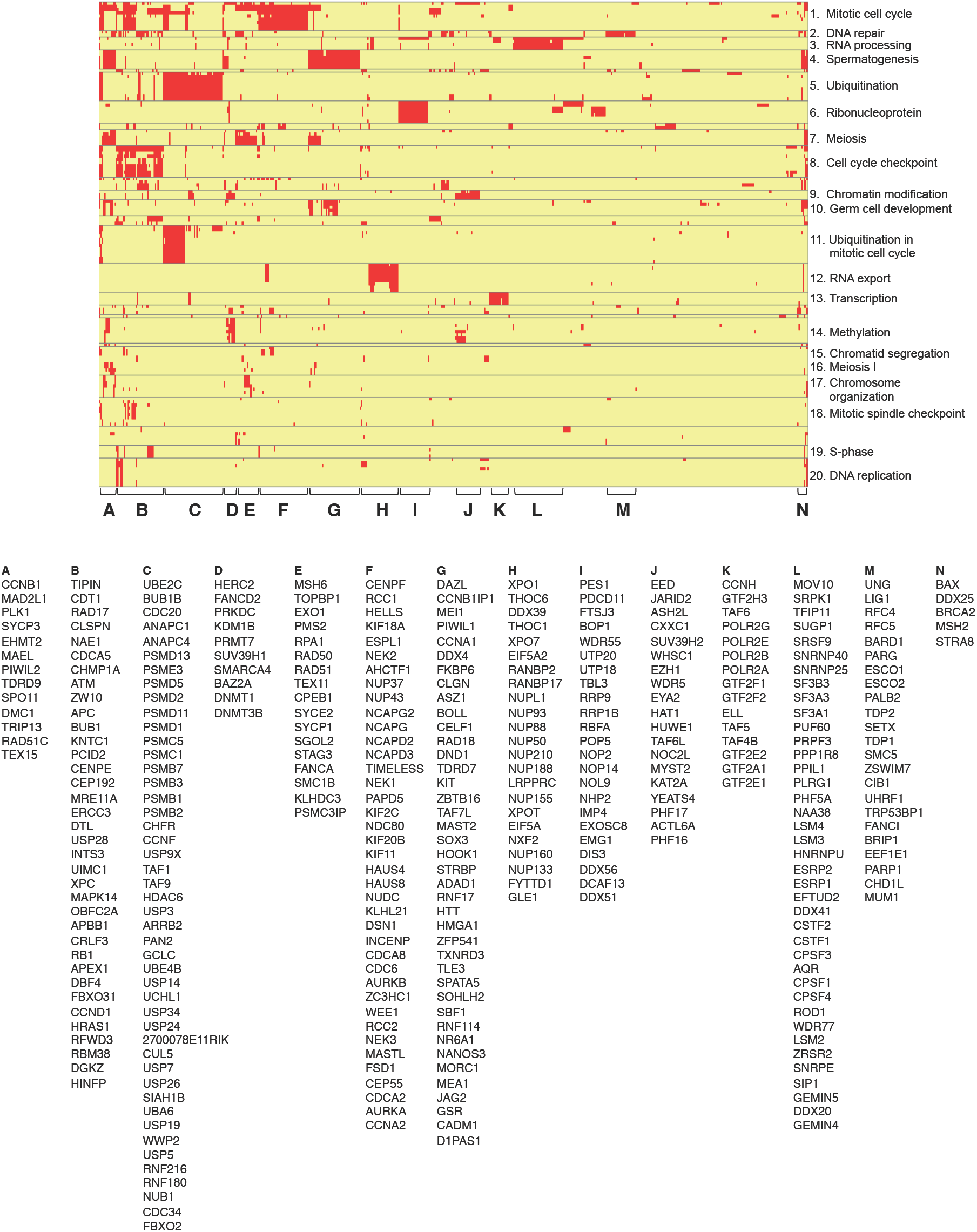
Hierarchical clustering of enriched GO terms (*P*<0.01) and genes associated with the set of 1462 genes that have decreased RNA levels in *DAZL* KO cells.

**Supplemental Figure 7.**
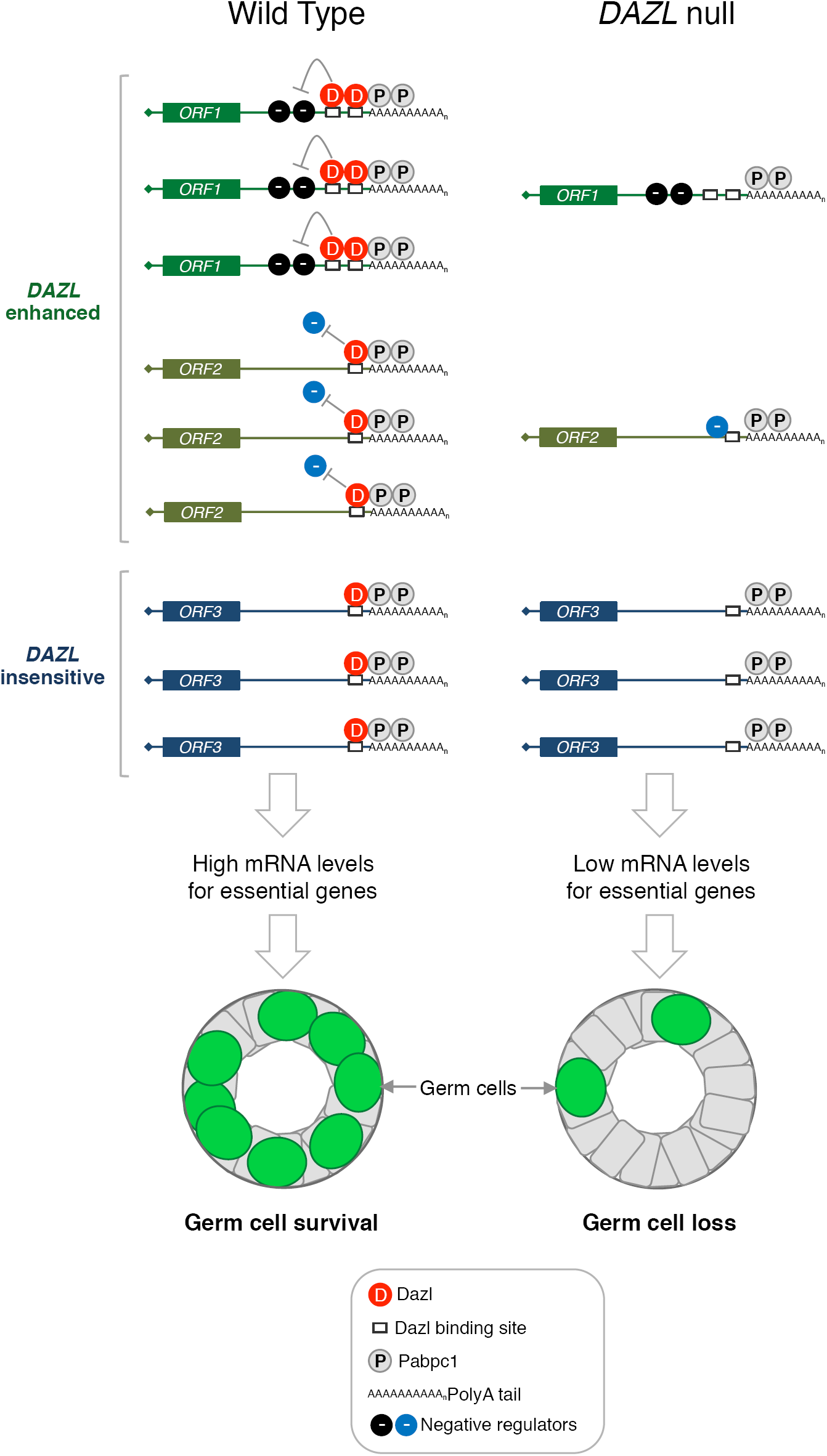
Proposed model for *DAZL*-dependent germ cell maintenance. Dazl binds a broad set of mRNAs via polyA-proximal GUU interactions facilitated by Pabpc1-polyA interactions. Dazl maintains high mRNA levels for select targets (*ORF1* and *ORF2*), potentially by blocking the function or binding of negative regulatory factors. In the absence of Dazl, *ORF1* and *ORF2* mRNAs are reduced, whereas *ORF3* mRNAs (not subject to negative regulation) are unaffected.

## Supplemental Tables

**Table S1.** Annotation of 11,297 BR3 clusters in RefSeq fragments. Clusters that overlap two different fragments in the same transcript (such as coding and UTR sequence) are denoted with “-“. Clusters that map to positions with alternative annotations (such as intron in one isoform and exon in another) are denoted with “/”.

| Genic location | BR3 Clusters | % BR3 | 5'UTR | 3'UTR | CDS | Intron |
| --- | --- | --- | --- | --- | --- | --- |
| 3UTR | 6397 | 70.59 |  | 6397 |  |  |
| 3UTR // 5UTR | 3 | 0.03 | 3 | 3 |  |  |
| 3UTR // CDS-3UTR | 10 | 0.11 |  | 10 | 10 |  |
| 3UTR // CDS-INTRON | 1 | 0.01 |  | 1 | 1 | 1 |
| 3UTR // INTRON | 157 | 1.73 |  | 157 |  | 157 |
| 3UTR // INTRON-3UTR | 5 | 0.06 |  | 5 |  | 5 |
| 3UTR // INTRON-CDS-3UTR | 3 | 0.03 |  | 3 | 3 | 3 |
| 3UTR-INTRON | 1 | 0.01 |  | 1 |  | 1 |
| 5UTR | 57 | 0.63 | 57 |  |  |  |
| 5UTR // CDS | 1 | 0.01 | 1 |  | 1 |  |
| 5UTR // INTRON | 7 | 0.08 | 7 |  |  | 7 |
| 5UTR-CDS | 37 | 0.41 | 37 |  | 37 |  |
| 5UTR-CDS // INTRON | 3 | 0.03 | 3 |  | 3 | 3 |
| 5UTR-CDS // INTRON-5UTR-CDS | 3 | 0.03 | 3 |  | 3 | 3 |
| 5UTR-CDS-INTRON | 3 | 0.03 | 3 |  | 3 | 3 |
| 5UTR-INTRON | 4 | 0.04 | 4 |  |  | 4 |
| CDS | 741 | 8.18 |  |  | 741 |  |
| CDS // 3UTR | 1 | 0.01 |  | 1 | 1 |  |
| CDS // INTRON | 16 | 0.18 |  |  | 16 | 16 |
| CDS // INTRON-CDS | 1 | 0.01 |  |  | 1 | 1 |
| CDS-3UTR | 775 | 8.55 |  | 775 | 775 |  |
| CDS-3UTR // CDS-3UTR-INTRON | 2 | 0.02 |  | 2 | 2 | 2 |
| CDS-3UTR // CDS-INTRON | 4 | 0.04 |  | 4 | 4 | 4 |
| CDS-3UTR // INTRON | 15 | 0.17 |  | 15 | 15 | 15 |
| CDS-3UTR-INTRON | 5 | 0.06 |  | 5 | 5 | 5 |
| CDS-INTRON | 226 | 2.49 |  |  | 226 | 226 |
| INTRON | 340 | 3.75 |  |  |  | 340 |
| INTRON-3UTR | 10 | 0.11 |  | 10 |  | 10 |
| INTRON-3UTR // INTRON-CDS-3UTR | 1 | 0.01 |  | 1 | 1 | 1 |
| INTRON-5UTR | 2 | 0.02 | 2 |  |  | 2 |
| INTRON-5UTR // 5UTR | 1 | 0.01 | 1 |  |  | 1 |
| INTRON-5UTR-CDS | 6 | 0.07 | 6 |  | 6 | 6 |
| INTRON-5UTR-CDS-INTRON | 3 | 0.03 | 3 |  | 3 | 3 |
| INTRON-5UTR-INTRON | 1 | 0.01 | 1 |  |  | 1 |
| INTRON-CDS | 126 | 1.39 |  |  | 126 | 126 |
| INTRON-CDS // 3UTR | 2 | 0.02 |  | 2 | 2 | 2 |
| INTRON-CDS // INTRON-CDS-3UTR | 1 | 0.01 |  | 1 | 1 | 1 |
| INTRON-CDS-3UTR | 62 | 0.68 |  | 62 | 62 | 62 |
| INTRON-CDS-3UTR-INTRON | 1 | 0.01 |  | 1 | 1 | 1 |
| INTRON-CDS-INTRON | 28 | 0.31 |  |  | 28 | 28 |

